# Structural remodelling of the carbon-phosphorus lyase machinery by a dual ABC ATPase

**DOI:** 10.1101/2022.06.09.495270

**Authors:** Søren K. Amstrup, Nicholas Sofos, Jesper L. Karlsen, Ragnhild B. Skjerning, Thomas Boesen, Jan J. Enghild, Bjarne Hove-Jensen, Ditlev E. Brodersen

## Abstract

Phosphorus is an essential macronutrient for all microorganisms and can be extracted from the environment by several metabolic pathways. In *Escherichia coli,* the 14-cistron *phn* operon encoding the carbon-phosphorus (C-P) lyase enzymatic machinery allows for extraction of phosphorus from a wide range of phosphonates characterised by the highly stable C-P bond.^1, 2^ As part of a complex, multi-step pathway, the PhnJ subunit was proposed to cleave the C-P bond via a radical reaction, however, the details of the mechanism were not immediately compatible with the structure of the 220 kDa PhnGHIJ C-P lyase core complex, leaving a significant gap in our understanding of phosphonate breakdown in bacteria.^3, 4^ Here we show using single-particle cryogenic-electron microscopy that PhnJ mediates binding of a unique double dimer of ATP-binding cassette (ABC) proteins, PhnK and PhnL to the core complex. ATP hydrolysis by PhnK induces drastic structural remodelling leading to opening of the core and reconfiguration of a metal-binding site located at the interface between the PhnI and PhnJ subunits. Our results offer new insights into the mechanism underlying C-P lyase and uncover a hitherto unknown configuration of ABCs that have wide-ranging implications for our understanding of the role of this module in biological systems.

---

Bacteria have evolved elaborate mechanisms to extract essential macronutrients, such as phosphorus, sulphur, nitrogen and carbon, from their environments. In the Gram-negative γ-proteobacterium, *Escherichia coli,* phosphate limitation induces Pho regulon expression, including the 14-cistron *phn* operon *(phnCDEFGHIJKLMNOP),* enabling uptake and breakdown of phosphonates as an alternative source of phosphorus.^2, 5, 6^ Phosphonates are structurally similar to phosphate esters, transition state analogues, and other primary metabolites but contain a highly stable carbonphosphorous (C-P) bond, which is resistant to hydrolytic cleavage.^7^ Phosphonate components are therefore highly valuable in biotechnology and agriculture, as detergents, herbicides (e.g. glyphosate/RoundUp®), and antibiotics.^8^ The *phn* operon encodes an ABC importer and periplasmic phosphonate binding protein *(phnCDE)* used for phosphonate uptake^9^, a transcriptional regulator (*phnF*)^10^, as well as the complete enzymatic machinery required for the degradation of phosphonates *(phnGHIJKLMNOP)* and incorporation of phosphorus into general metabolism via 5-phospho-*α*-*D*-ribosyl-1-diphosphate (PRPP)^11, 12, 13, 14^ The C-P lyase pathway has a remarkably wide substrate spectrum and is capable of handling aliphatic as well as aromatic phosphonates.^13, 15, 16^ For this reason, and due to the many applications of phosphonates, there is a strong motivation to understand the mechanisms of phosphonate breakdown by bacteria. Detailed functional and mechanistic studies of individual enzymatic subunits encoded by the *phn* operon led to a proposed reaction mechanism in which the phosphonate moiety is initially coupled to the 1’ position of either ATP or GTP while displacing the nucleobase in a reaction catalysed by PhnI and requiring the presence of PhnG, PhnH, and PhnL.^11^ PhnM was shown to then release pyrophosphate from the resulting 5’-phosphoribosyl-*α*-1 -phosphonate to allow for PhnJ to cleave the C-P bond via a strictly anaerobic *S*-adenosyl methionine (SAM)-dependent radical reaction mechanism. Finally, the combined action of PhnP and PhnN converts the resulting cyclic ribose into PRPP (**Supplementary Figure 1**).^12, 13^

The complexity of the reaction, the large number of protein subunits involved, and anaerobic requirement has meant that it has been challenging to obtain a complete, mechanistic understanding of the C-P lyase pathway. The crystal structure of a core enzyme complex, Phn(GHIJ)_2_, revealed a symmetrical heterooctamer consisting of a compact central tetramer of PhnI and PhnG subunits interacting with more distal copies of PhnJ and PhnH.^4^ An intriguing feature of this structure is that the site of the Fe_4_S_4_ cluster, required for radical generation and enzyme activation, is located at a significant distance (30 Å) from Gly32 of PhnJ, which had been shown by deuterium exchange studies to receive a hydrogen radical through exchange with activated SAM near the Fe_4_S_4_ site.^3^ The structure also revealed another putative functional site at the interface of PhnI and PhnJ containing a Zn^2+^ coordinated by two conserved His residues (PhnI His328, His333), both found to be essential for growth on phosphonate. Finally, it was found that of the two non-transporter ABC proteins encoded by the *phn* operon, PhnK and PhnL, a single subunit of PhnK can bind to PhnJ via a conserved helical region, the central insertion domain (CID).^4^ ABC modules are associated with metabolite transporters where they interact as dimers with a transmembrane domain to bind and hydrolyse ATP, thus generating the energy required for the transport process.^17^ Therefore, the structure of the C-P lyase bound to a single PhnK subunit was surprising and did not reveal how this subunit would function in the catalytic process, nor could it suggest a role for the other ABC protein, PhnL.

## RESULTS

### A single PhnK subunit binds to the C-P lyase core complex in a disordered state not compatible with ATP hydrolysis

To better understand the interaction of ABC subunits with C-P lyase and the role of these modules in catalysis, we initially purified the PhnK-bound core complex from a plasmid encoding *phnGHIJK* as previously described^4^ and determined a high-resolution structure by cryogenic-electron microscopy (cryo-EM). An initial map generated using 901,800 particles (**Supplementary Figure 2**) resulted in a structure at an overall resolution of 2.2 Å (using the gold standard Fourier shell correlation (FSC) 0.143 cut-off criterion^18^) of the C-P lyase core complex bound to a single PhnK subunit as previously observed^4, 17^ (**Figure 1a**). However, PhnK displayed a significantly lower resolution than the core complex, so we used 3D classification with signal subtraction to focus the refinement on PhnK and obtained a final class consisting of 50,323 particles at a resolution of 2.6 Å. This map displayed improved density for PhnK and allowed for tracing of the complete fold of the protein (**Figure 1a, Supplementary Figure 2 and Supplementary Table 1**). Variability analysis revealed several distinct conformations of PhnK corresponding to a hingelike motion of the ABC domain around the site of interaction (**Figure 1b**).^19, 20^ The binding of PhnK to the Central Insertion Domain (CID) of PhnJ is driven primarily by electrostatic interactions involving negatively charged residues in PhnJ (Glu149, Asp226, Asp228, and Asp229) and a positive patch on PhnK (Arg 78, Arg82, and Arg116) (**Figure 1c and 1d**). These interactions are further stabilised by hydrophobic interaction between PhnJ Tyr 158 and PhnK Tyr 118.

**Figure 1.**
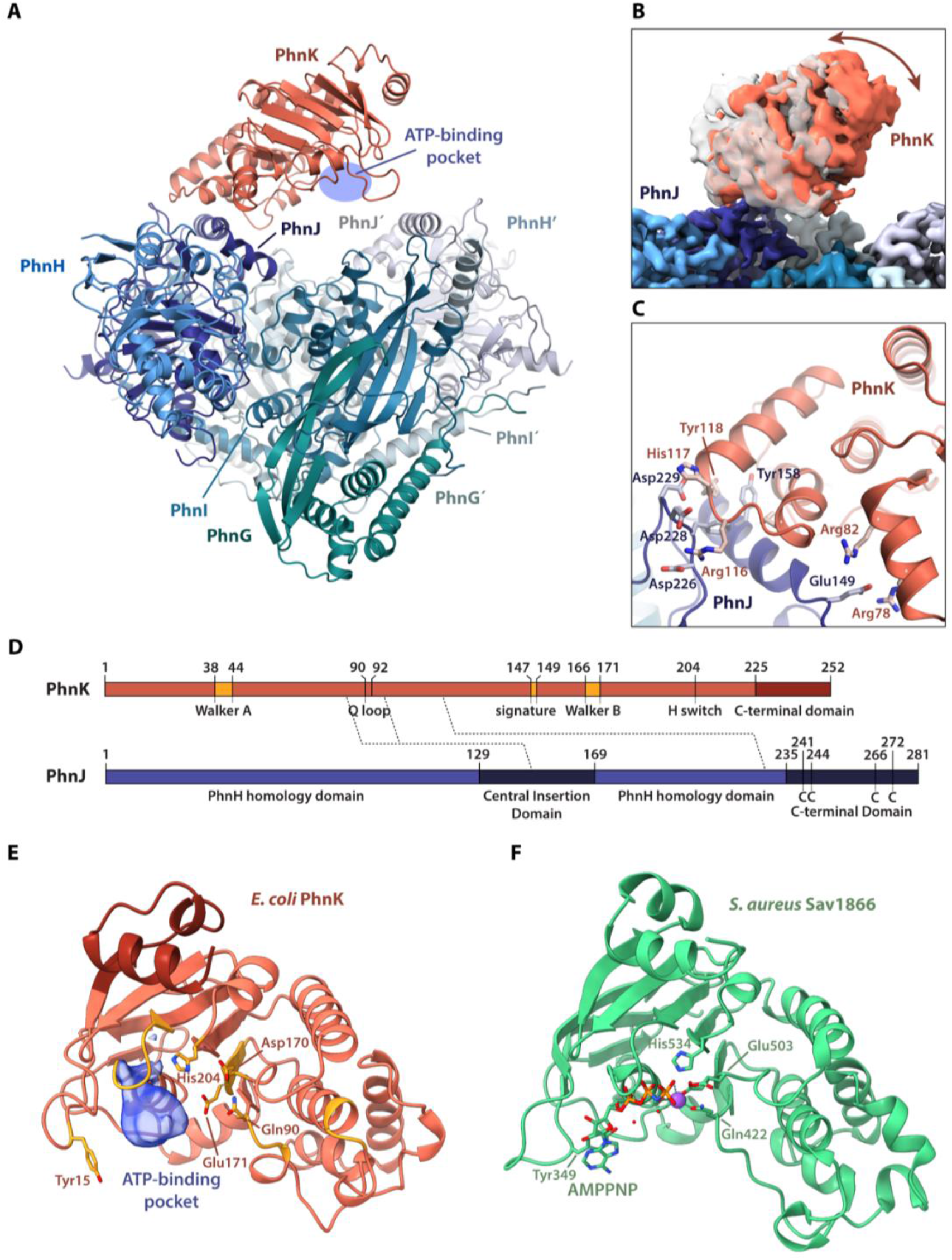
The C-P lyase core complex can bind a single, flexible PhnK subunit. **a,** Overview of the structure of the C-P lyase core complex (PhnGHIJ, blue/green) bound to a single subunit of PhnK (red) with the names of individual subunits indicated. The ATP-binding pocket of PhnK is shown with a blue shape. **b,** surface representation showing the extent of tilting motion observed in PhnK. The two extreme states are shown in white and red, respectively. **c,** Close-up of the interaction between PhnK (red) and the PhnJ central insertion domain (CID, blue) with relevant residues shown as labelled sticks. **d,** Overview of the domain structure of PhnK (top, red) and PhnJ (bottom, blue) with residue numbers indicated. For the NBD protein PhnK the location of the core ABC motifs (Walker A and B, Q loop, signature motif, and H switch) are indicated in orange and the C-terminal domain in dark red. For PhnJ, the unique features that distinguish the protein from PhnH with which it is homologous (the Central Insertion Domain, CID, residues 129-169, and the C-terminal Domain, CTD, residues 235-281) are shown in darker blue. The four Cys residues near the C-terminus involved in Fe_4_S_4_ cluster binding are indicated with “C”. Finally, interactions between the two proteins are shown with dashed lines. **e,** Overview of the PhnK structure from the Phn(GHIJ)_2_K structure with catalytically important residues shown as sticks. An extra density observed in several 3D classes at the nucleotide binding site is shown in blue. **f,** Structure of a single NBD of the *S. aureus* Sav1866 multidrug transporter (PDB ID: 2ONJ)^25^ shown in the same orientation as PhnK with the nucleotide (AMPPNP) as sticks and the Mg^2+^ ion as a purple sphere.

PhnK adopts a classical nucleotide-binding domain (NBD) fold like those found in ABC transporters and all the required catalytic motifs (the A/P/D/Q loops, H switch, Walker A/B, and ABC signature motifs, **Figure 1d and 1e**) are conserved. Of these, the A, P, and D loops constitute a highly important region directly involved in ATP binding and hydrolysis.^21^ In a subset of 3D classes, extra density was visible near the P-loop of PhnK, suggesting the presence of a nucleotide (**Figure 1e and Supplementary Figure 3)**. The density overlaps partly with that of ATP as bound to a dimer of NBD modules in ABC transporters (**Figure 1f**), but the lack of high-resolution features for this density suggests that binding is disordered and consequently, that PhnK is not in a catalytically competent state. This is consistent with the general understanding that ABC modules are only active in the dimeric state. In summary, we conclude that the structure of a single PhnK subunit bound to the C-P lyase core complex suggests binding of the NBD in this state is disordered, can potentially bind nucleotide, but not engage in ATP hydrolysis. Moreover, and in contrast to previous observations^17^, we observe no structural changes in the core complex upon binding of a single subunit PhnK that could explain the functional discrepancies presented by the structure of the C-P lyase core complex.

### PhnK and PhnL associate as a unique double dimer to the C-P lyase core complex

To capture PhnK in a stable, nucleotide bound conformation, we introduced the E171Q mutation in the Walker B motif (**Figure 1e**), which is known to allow binding, but not hydrolysis of ATP in ABC modules^22^. We then purified the C-P lyase complex as expressed from a plasmid encoding the full C-P lyase enzymatic machinery *(phnGHIJKLMNOP)* using a C-terminal double Strep tag on PhnK and in the presence of the non-hydrolysable ATP analogue, β,γ-imidoadenosine 5’-triphosphate (AMPPNP). Structure determination of this sample by cryo-EM single-particle analysis at 2.1 Å resolution (59,737 particles, C2 symmetry, **Supplementary Figure 4 and Supplementary Table 1**) revealed the core complex bound to a dimer of PhnK in a conformation reminiscent of the ATP-bound state of ABC transporters (**Figure 2, red subunits**). At the interface of the two PhnK subunits, clear density for AMPPNP allowed for modelling of a nucleotide in both active site pockets (**Figure 2a and Supplementary Figure 5a, left**). Inside PhnK, the adenosine nucleotide binds in a way very similar to what has been observed for other ABC modules (**Supplementary Figure 5b**). Despite the presence of an additional PhnK subunit and the tight interaction between the two NBDs, very few structural changes are observed in the C-P lyase core complex compared to the unbound state (**Supplementary Figure 5c**). Remarkably, the structure also revealed two copies of an additional protein situated on top of the PhnK subunits, which we could confirm by mass spectrometry as well as inspection of the EM density to be PhnL (**Figure 2, yellow subunits, and Supplementary Figure 6a**). Like PhnK, PhnL has an NBD fold like those found in ABC transporters but differs by lacking a conserved C-terminal domain and by the presence of a long β hairpin near the N terminus that results from extension of β1 and β2 (**Figure 2**). The association of a PhnL subunit with each of the two PhnK subunits thus generates a unique double dimer of ABC subunits. In this hetero-dodecameric, 327 kDa enzyme complex, PhnL is in an open *apo* state without visible density for a nucleotide. The interaction between PhnK and PhnL is mediated by a wide range of specific interactions, mostly via the C terminal domain of PhnK, which contains two α helices (α8 and α9) that facilitate most contacts with PhnL, including the extended β hairpin (**Figure 2b-d**). The interactions are primarily ionic with the binding region (top side) on PhnK having an overall negative charge (residues 231-247) while PhnL (underside) is positive (**Supplementary Figure 6b)**. In addition, π-π stacking is observed between PhnK Trp29 and PhnL Arg82 (**Figure 2b**). The Arg in PhnL is fully conserved in orthologues, while the Trp in PhnK is functionally conserved for its stacking ability (**Supplementary Figure 6c)**. In summary, we conclude that both non-transporter ABC subunits encoded by the *phn* operon, PhnK and PhnL, can bind simultaneously to the C-P lyase core complex in a unique dual dimer conformation with PhnK in the ATP-bound state. To our knowledge, this type of interaction between two ABC dimers has not been observed before and might therefore represent a new functional mode of nucleotide-binding domains.

**Figure 2.**
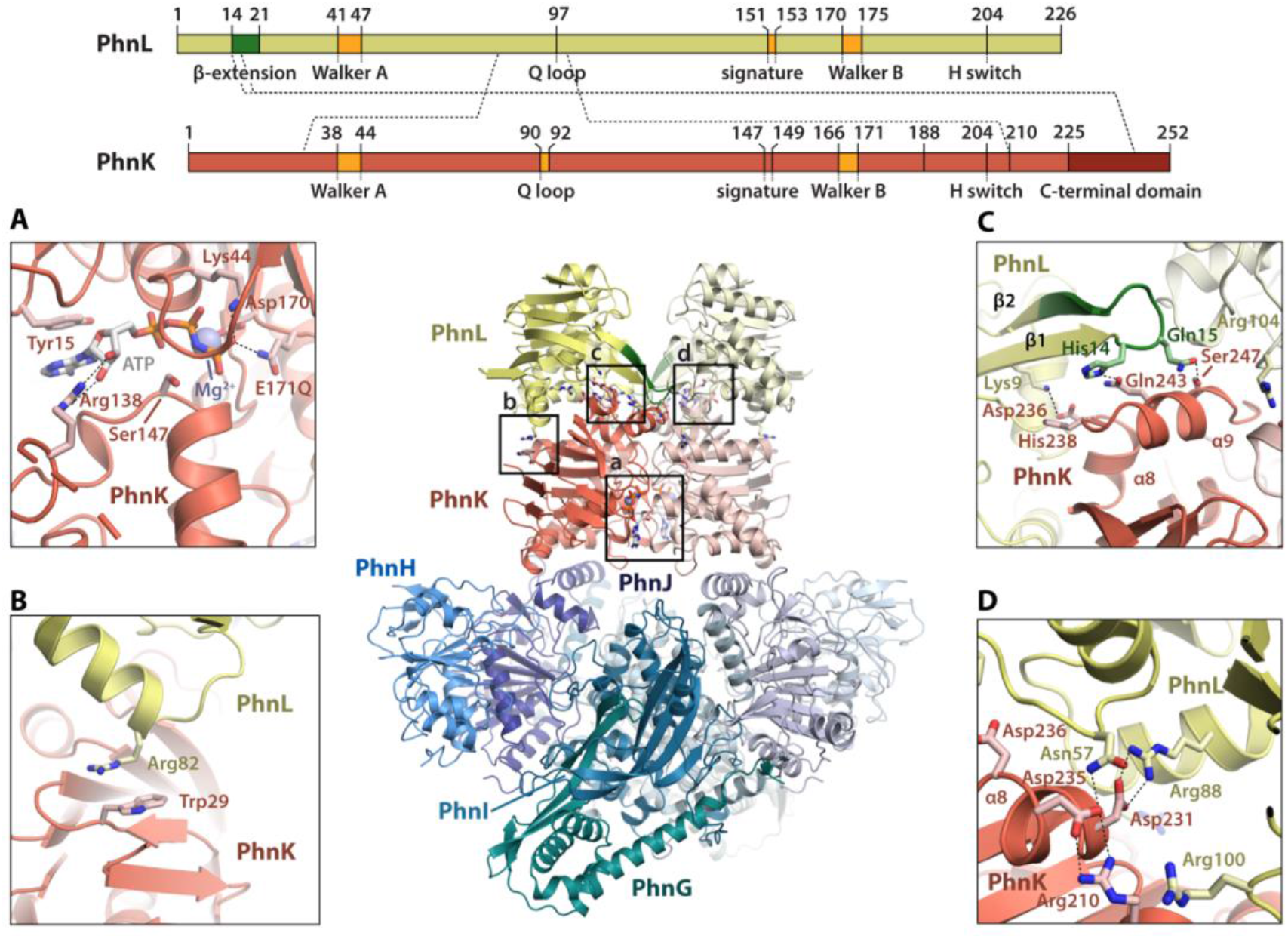
PhnGHIJKL displays a novel ABC dimer sandwich interaction. Top, an overview of the domain structure and sequences features of PhnL (top, yellow) and PhnK (middle, red) with residue numbers indicated. For the ABC proteins, the location of the core ATPase motifs (Walker A and B, Q loop, signature motif, and H switch) are indicated in orange. In PhnL, the unique β hairpin extension (residues 7-14) is shown in green and for PhnK, the C-terminal domain is shown in red. Finally, interactions between proteins are shown with dashed lines. Centre, an overview of the structure of PhnGHIJKL as cartoon with the C-P lyase core complex PhnGHIJ in shades of blue/green and corresponding light colours for the other half, the PhnK dimer in red/light red, and the PhnL dimer in yellow/light yellow. Boxes indicate the location of the close-up views a-d. **a,** Details of the PhnK ATP binding site with the nucleotide (white) shown alongside relevant, interacting side chains (labelled sticks), and the Mg^2+^ ion as a blue sphere. **b,** Stacking interaction observed between PhnK Trp29 and PhnL Arg82. **c,** Interaction between the PhnK C-terminus (helices α8 and α9, residues 236247) and the PhnL β hairpin extension (residues 7-14). **d,** Charged interactions observed between the PhnK C-terminal region (residues 210-236) and PhnL.

### Binding and activity of both PhnK and PhnL are essential for phosphonate utilisation *in vivo*

To probe the importance of the specific interactions between PhnK and the core complex as well as the functional integrity of PhnK and PhnL for phosphonate breakdown, we used genetic complementation to test if alteration of specific residues would affect the ability of *E. coli* to grow on phosphonate as a sole source of phosphorus (**Figure 3a**). For this experiment, we transformed the *E. coli Δphn* strain HO1488 with a plasmid-borne copy of the *phn* operon (pSKA03, see Methods for details), either containing wild type or mutated variants of individual proteins. Initially, we could show that the inability of the HO1488 strain to grow on either methylphosphonate or 2-aminoethylphosphonate could be genetically complemented by introduction of the pSKA03 plasmid (**Figure 3a**). In contrast, disruption of residues at the PhnK-PhnJ interaction interface (PhnJ E149A, Y158A or PhnK R78A/R82A) or inhibition of ATPase activity of either PhnK or PhnL by mutation of the catalytically active glutamate (PhnK E171Q or PhnL E175Q)^22^ abolished growth on phosphonate as a sole source of phosphorus. Together, this demonstrates that both binding of PhnK to the core complex and activity of PhnK and PhnL are required for phosphonate breakdown *in vivo.*

**Figure 3.**
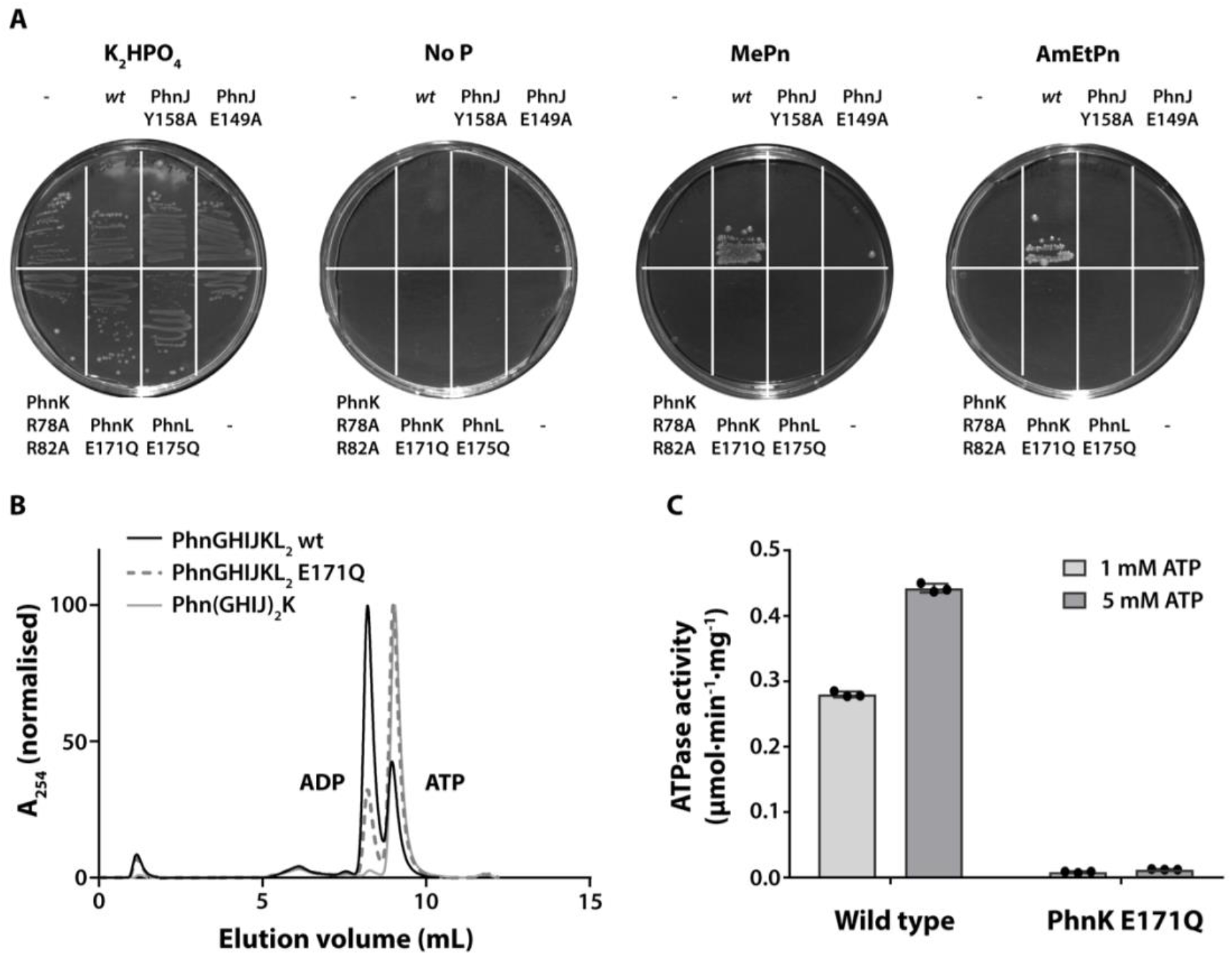
ATP hydrolysis by PhnK and PhnL is required for phosphonate utilisation *in vivo*. **a,** *In vivo* functional assay using the *E. coli* HO1488 strain *(ΔphnHIJKLMNOP)* either without plasmid (“-”) or complemented with pSKA03 containing the entire *phn* operon including the *pho* box and including the following specific mutations: wt (none), PhnJ Y158A, PhnJ E149A, PhnK R78A/R82A, PhnK E171Q, and PhnL E175Q. Cells were plated on MOPS minimal agar plates^26^ containing either K_2_HPO_4_, 2-AmEtPn, MePn or no added phosphate source (“no P”) as indicated. The plates are representative of two repetitions. **b,** ATP turnover catalysed by Phn(GHIJ)_2_K, Phn(GHIJKL)_2_ PhnK E171Q, and Phn(GHIJKL)_2_ WT, as indicated, measured by separation of nucleotide species by ion-exchange chromatography after overnight incubation with ATP. **c,** ATPase activity (μmol/min/mg) measured via a coupled assay in the presence of either 1- or 5-mM ATP for wild type PhnGHIJKL complex and PhnK W171Q complex, as indicated. The bars indicate the mean of the results from three independent reactions using the same protein stocks (dots) and error bars show standard deviation from the mean.

Since ATP hydrolysis by PhnK and PhnL thus is essential for the ability of *E. coli* to grow on phosphonate, we next asked if the protein complex consisting of the C-P lyase core complex bound to PhnK and PhnL, Phn(GHIJKL)_2_, is active in ATP hydrolysis *in vitro.* To answer this, the complex was purified both as wild type and in context of the PhnK E171Q ATPase mutation (**Supplementary Figure 7a and 7b**) and analysed for ATPase activity by two methods, incubation of ATP and separation of reaction products using ion exchange chromatography (**Figure 3b**) or a coupled enzymatic ATPase assay (**Figure 3c**). Both experiments revealed a solid ATPase activity for the wild type complex, which is almost absent in the PhnK E171Q mutant, suggesting that only PhnK and not PhnL can hydrolyse ATP under these conditions. In summary, we conclude that both PhnK and PhnL are required to be catalytically active to support phosphonate utilisation *in vivo* but that only PhnK displays ATPase activity in the purified complex.

### ATP turnover by PhnK induces large-scale conformational changes in the C-P lyase core complex

To capture potential, additional conformational states of the complex resulting from the ATPase activity, we next purified wild type Phn(GHIJKL)_2_ complex in the presence of ATP and EDTA (to initially prevent ATP hydrolysis by PhnK), then added Mg^2+^ to initiate the reaction immediately before spotting onto cryo-EM grids and plunge freezing. Cryo-EM data collection and processing under these conditions yielded several structures including a high-resolution structure of the Phn(GHIJKL)_2_ complex extending to 1.9 Å using a final set of 222,056 particles and imposing C2 symmetry (**Supplementary Figure 8 and 9, and Supplementary Table 1**). The conformation of this complex is essentially identical to the structure with PhnK bound to AMPPNP, however, inspection of the PhnK active site reveals that it contains ADP and P_i_ and consequently is an ATP post-hydrolysis state (**Supplementary Figure 5a, middle**). In this state, we also observe binding of ATP to both nucleotide binding sites in PhnL, even though this dimer is in the open (ADP) state (**Supplementary Figure 10a**). We believe that this is likely a consequence of the higher nucleotide concentration used to prepare this sample.

3D classification of the data collected under ATP turnover conditions yielded two additional structures of the core complex bound to two subunits of PhnK but not always PhnL (**Figure 4a, Supplementary Figure 8 and 9, and Supplementary Table 1**). In both structures, the density for PhnK is significantly poorer than in the stable PhnL-bound state, suggesting it is more flexible and possibly that PhnL acts to stabilise the closed conformation of PhnK (**Supplementary Figure 10b**). In one of these conformations (the “closed” state, 2.0 Å, 81,605 particles, C2 symmetry), the PhnK dimer is intact and resembles that found in the Phn(GHIJKL)_2_ structures, and inspection of the active site reveals clear density for both ATP and Mg^2+^ suggesting that this is a pre-hydrolytic ATP-bound state (**Figure 4a, closed, and Supplementary Figure 5a, right**). In this state, there is no evidence of PhnL in neither 3D density nor 2D class averages (**Supplementary Figure 10c and 10d**). In the other structure (the “open” state, 2.6 Å, 31.280 particles, C1 symmetry), the two PhnK subunits have separated in such a way that one subunit has moved about 25 Å away while the other has stayed in the position corresponding to the closed state (**Figure 4a, open**). Due to the tethering of PhnK to PhnJ, this movement also results in pulling one set of PhnJ and PhnH subunits away from the rest of the core complex and in the process exposing the Zn^2+^-binding site located at the PhnI-PhnJ interface (**Figure 4a**). Both PhnK subunits appear very flexible in this state (similar to the Phn(GHIJ)_2_K structure) and display a significantly lower local resolution than the rest of the complex. PhnL is not visible in the final map, but inspection of the 2D class averages reveals that PhnL is bound in a closed (possibly ATP) state (**Supplementary Figure 10c and 10d**).

**Figure 4.**
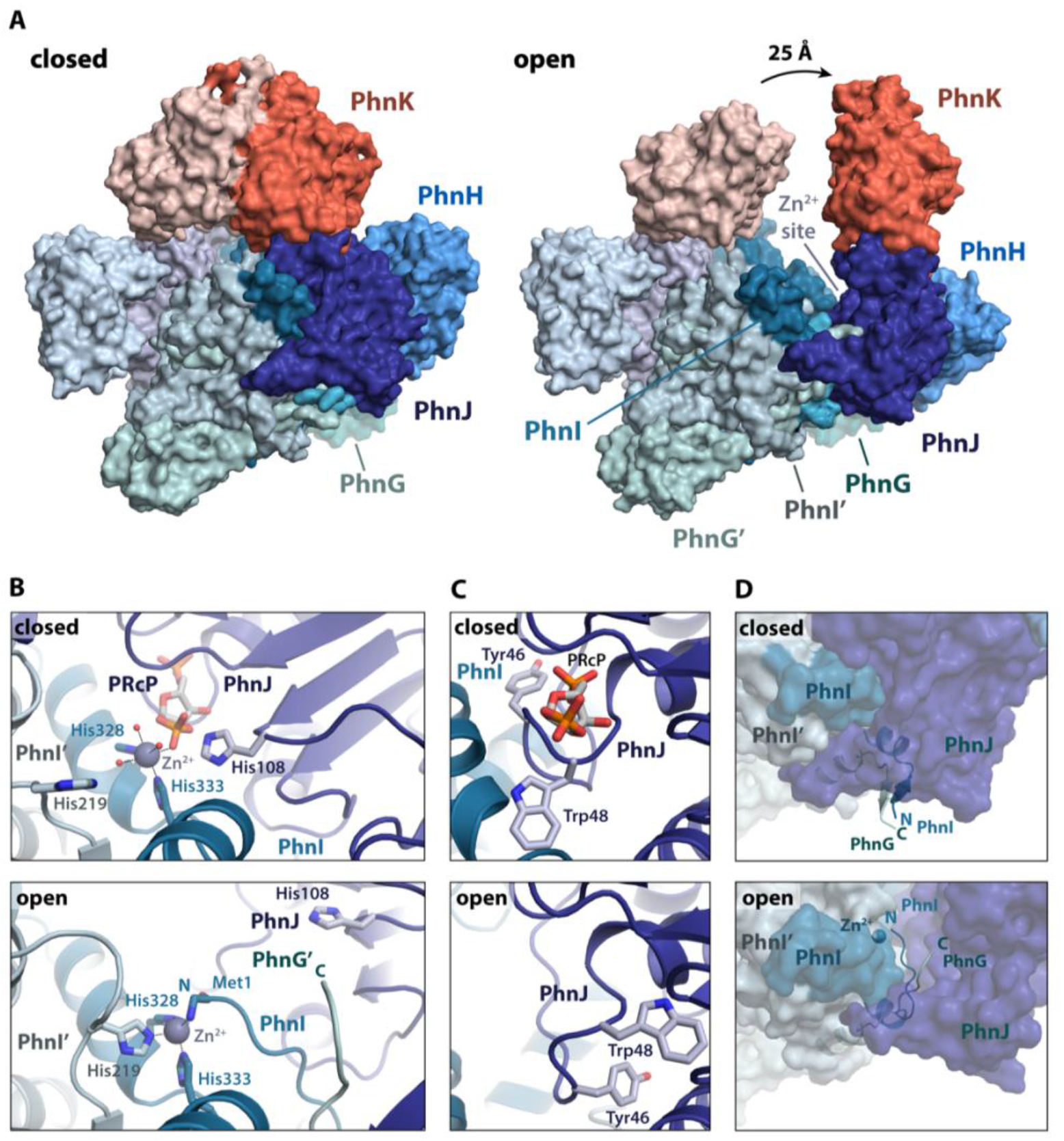
ATP hydrolysis by PhnK leads to opening of the C-P lyase core complex. **a,** Overview of the overall structural changes taking place in the Phn(GHIJK)_2_ complex between the closed (left) and open (right) states. The Phn(GHIJK)_2_ complex is shown as a surface representation with the C-P lyase core complex in shades of blue/green and PhnK in red/light red. **b,** Details of the active site at the interface between PhnI and PhnJ in the closed (top) and open (bottom) states. The Zn^2+^ ion is shown as a grey sphere with coordination geometry indicated and relevant, interacting residues with labelled sticks. **c,** Details of the active site pocket with bound PRcP as found in the closed (top) and without in open (bottom) states. **d,** Coordinated movement of the PhnI N terminus and PhnG C terminus from the surface of the PhnGHIJK complex in the closed state (top) to the active site in the open state (bottom).

Given that the core complex has remained stable in all previous structures, the concomitant movement of PhnK, PhnJ, and PhnH in the open state represents a dramatic architectural rearrangement with significant functional implications. In the closed state, both PhnI subunits and one PhnJ subunit interact tightly to form a Zn^2+^-binding site located at the interface of the three proteins (the “His site”)^4^. In addition to two histidine residues originating from PhnI (His328, His333), the Zn^2+^ ion is coordinated by three water molecules and an oxygen from an unknown ligand, which was also observed previously but could not be modelled, in an octahedral arrangement (**Figure 4b, “closed” and Supplementary Figure 11a)**.^4^ Based on inspection of the EM density we were able to model the ligand as 5-phospho-*α*-*D*-ribose-1,2-cyclic-phosphate (PRcP) an intermediate of the C-P lyase reaction (**Supplementary Figure 1 and 11b**).^11, 13^ Since this compound was not added to the protein sample before structure determination, we believe it was carried along during purification and could have been formed in a side reaction during overexpression in the absence of phosphonate. The ligand is tightly bound at this site with specific interactions between the ribose moiety and PhnJ Arg107 and Gln124, contacts between the 5’ phosphate and a loop (residues 47-50) in PhnJ and finally between two oxygen atoms of the 1,2-cyclic phosphate, PhnJ His108, and Zn^2+^ (**Supplementary Figure 11b**). We note that there is a significant cavity next to what would correspond to the phosphonate atom of a natural substrate, which could explain the wide substrate specificity of C-P lyase (**Supplementary Figure 11c**).

On the contrary, no ligand density is observed in the Zn^2+^ site in the open state, which has undergone a dramatic rearrangement, whereby the loop interacting with the 5’ phosphate of the ligand in the closed state has been completely remodelled (**Figure 4c**), allowing His219 from a nearby PhnI chain and the N-terminal NH_2_ group of PhnI to complete the Zn^2+^ coordination sphere, which is now tetrahedral (**Figure 4b, “open” and Supplementary Figure 11d)**. Meanwhile, the PhnJ His108, which interacts with the 1,2-cyclic phosphate group of PRcP in the closed state has moved ~11 Å away from the metal ion. Intriguingly, the extended C terminus of PhnG, which in the closed state is located on the outside of the C-P lyase core complex where it forms tight interactions with the N terminus of PhnI using an antiparallel β sheet structure, is carried along with the PhnI N terminus as it relocates to the Zn^2+^ site (**Figure 4d**). Together, the PhnI and PhnG termini, therefore both undergo dramatic movements, in both cases more than 30 Å, a structural change that likely has several important, functional implications. We also note that the involvement of both chains of PhnI in Zn^2+^ coordination in the open state elegantly explains, along with the symmetry requirement for binding an ABC dimer, why the C-P lyase core complex must be a dimer of PhnGHIJ, thus solving a longstanding conundrum concerning the reason for the higher-order structure of C-P lyase.

## DISCUSSION

In this paper, we present structures of several states of the *E. coli* C-P lyase core complex bound to the ABC subunit PhnK showing that neither PhnK binding nor ATP binding to a dimer of PhnK *per se* alters the conformation of the core complex. This is in stark contrast to the classical role of NBDs in ABC transporters, where binding of ATP induces a more compact dimer and concomitant changes in the transmembrane segment, leading to eversion as part of the transport process.^23^ Thus, it appears that the stable ground state of the C-P lyase core complex corresponds to the ATP-bound conformation of PhnK. Moreover, we show that binding of a dimer of PhnK in the ATP-bound state to the C-P lyase core complex allows for association of a similar dimer of the homologous ABC protein, PhnL. This observation raises several questions regarding activation of the ATPase activities of both PhnK and PhnL, such as whether ATP hydrolysis happens simultaneously or sequentially in the two proteins. As a starting point, our *in vivo* complementation assays clearly show that ATPase activity of both PhnK and PhnL is required for phosphonate utilisation and thus, catalysis *in vivo.* But on the other hand, only PhnK appears to be active in ATP hydrolysis *in vitro*. We believe that a possible explanation for this discrepancy is the presence of the lengthy purification tag on PhnK, which in the Phn(GHIJKL)_2_ complex is located between the two PhnL subunits and thus likely preventing ATP binding and full closure of the second dimer, at least when PhnK is in the ATP-bound state (**Supplementary Figure 11e**). Structure determination under ATP turnover conditions allowed us to capture a post-hydrolysis structure for PhnK as bound to ADP + P_i_. For the ABC transporters, this conformation is difficult to capture due to fast release of P_i_ and energy upon ATP hydrolysis^24^, so this might suggest that, in the case of C-P lyase, PhnL serves to control phosphate release from PhnK and therefore potentially also provides an explanation for the requirement for two types of ABC modules. It then raises the question of how the ATPase activity of PhnL is controlled, however, which remains to be answered.

Both PhnK and PhnL bind in the same orientation to their partners, the C-P lyase core complex and PhnK, respectively, as the corresponding NBDs bind the transmembrane domains in the ABC transporters, but the structure of the target binding region differs in both cases from the typical coupling helix found in ABC transporters (**Figure 5**). In both cases, interaction between the NBD and its partner uses another binding geometry and appears to require unique elements, such as the extended β hairpin in PhnL and the C terminal domain of PhnK (**Figure 2, top and Figure 5**). Interestingly, the C terminal domain is found in other ABC proteins, such as the *Staphylococcus aureus* multidrug transporter, Sav1866 (**Figure 5a**)^25^, and is in fact a general feature of ABC modules.

**Figure 5.**
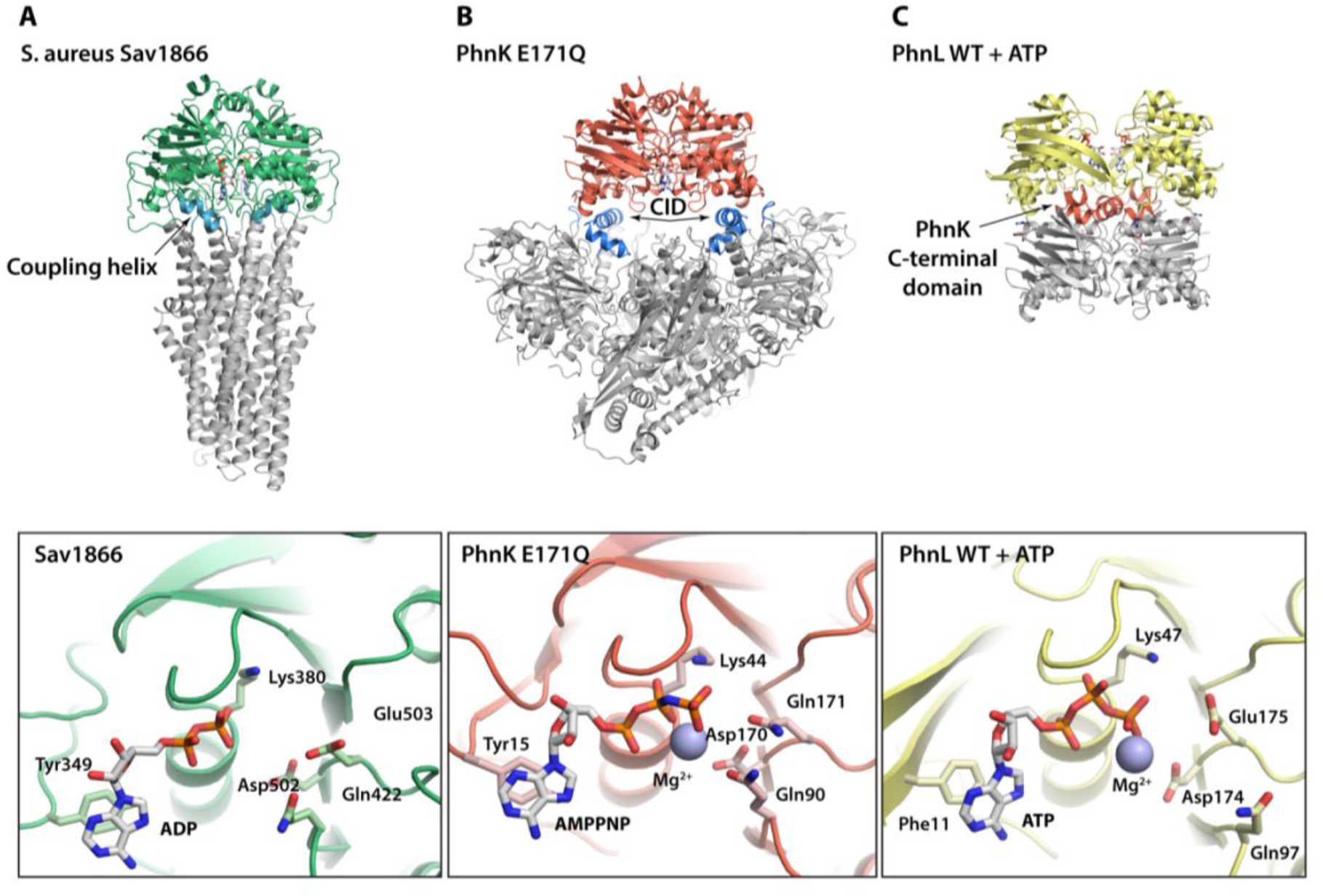
Comparison to the ABC transporters. **a,** Overview of the structure of the *S. aureus* Sav1866 ABC transporter in the ADP-bound conformation (top) with the ABC module in green and transmembrane domain in grey, except for the coupling helix, responsible for the interaction between the two, which is shown in cyan. The bottom panel shows details of the ATP binding site (bound to ADP). **b,** Overview of the structure of the C-P lyase core complex with a dimer of PhnK E171Q in the ATP bound conformation (top) with the PhnK dimer in red and the C-P lyase core complex in grey except for the PhnJ Central Insertion Domain (CID), responsible for PhnK binding, which is shown in blue. The bottom panel shows the details of the PhnK ATP binding site (bound to AMPPNP). **c,** Overview of the structure of the PhnK ABC dimer bound to PhnL in the open conformation (top) with the PhnL dimer in yellow and PhnK in grey except for the C-terminal region, responsible for the interaction, which is shown in red. The bottom panel shows details of ATP bound to the PhnL active site.

In any case, the double dimer of PhnK and PhnL observed in C-P lyase is novel and significantly expands the known ways in which NBD modules can bind their partner proteins. It also indicates that the overall fold of the binding partner is of less importance for interaction than specific interactions at the interface. As hypothesised above, PhnL could potentially function to inhibit P_i_ release from PhnK post hydrolysis and thus control when the energy generated from ATP hydrolysis is released, likely to break open the core complex as shown. The EM density suggests that this C-terminal region of PhnK becomes more ordered in the presence of PhnL. This effect extends towards helix 6 and the D loop, which could act as a gate to control P_i_ release. In this model, stabilisation of the PhnK D loop by PhnL would prevent P_i_ release.

Given that the C-P lyase core complex has appeared monolithic in all previous structures, we believe that the dramatic structural rearrangement from a closed to an open conformation is highly significant and will provide new ways of understanding the complex catalytic process of phosphonate breakdown by C-P lyase. Presumably, the inaccessible nature of the metal binding site between PhnI and PhnJ prevents ligand dissociation in the closed state. On the flip side, this suggests that opening of the complex and full occupation of the Zn^2+^ would happen along with exchange of the bound ligand. We therefore propose that the role of opening of the core complex by PhnK is important for substrate exchange. The reason for maintaining such a tightly controlled and energy-demanding substrate exchange mechanism, could be to control the local chemical environment required to perform the radical driven C-P lyase mechanism. Likewise, substrate binding might be a prerequisite for closure of the stable C-P lyase core complex, which would explain why there is always density for a ligand near the Zn^2+^ in the closed state.^4^ In summary, the structures of the C-P lyase complex presented here allow us to propose a model for a part of the pathway (**Figure 6**). According to this model, interaction of PhnK in the ATP-bound conformation with the C-P lyase core complex already bound to 5-phospho-*α*-*D*-ribose-1-phosphonate at the Zn^2+^ site resulting from the first two steps of the pathway (**Supplementary Figure 1**), allows for ATP binding in PhnL as well leading to a closed core complex that might signal to PhnJ to proceed with cleavage of the C-P bond, possibly at the identified Zn^2+^ binding site. Subsequently, ATP hydrolysis in PhnK would lead to opening of the core complex. Our 2D analysis (**Supplementary Figure 10c and 10d**) then suggests that this could happen concomitantly with association of the PhnL subunits and likely, substrate exchange. The product, PRcP, would then diffuse out for further processing by PhnP and PhnN allowing a new substrate molecule to bind. Renewed closure of the core complex could be the signal to trigger ATP hydrolysis in PhnL causing the subunits to dissociate and ADP + P_i_ to leave.

**Figure 6.**
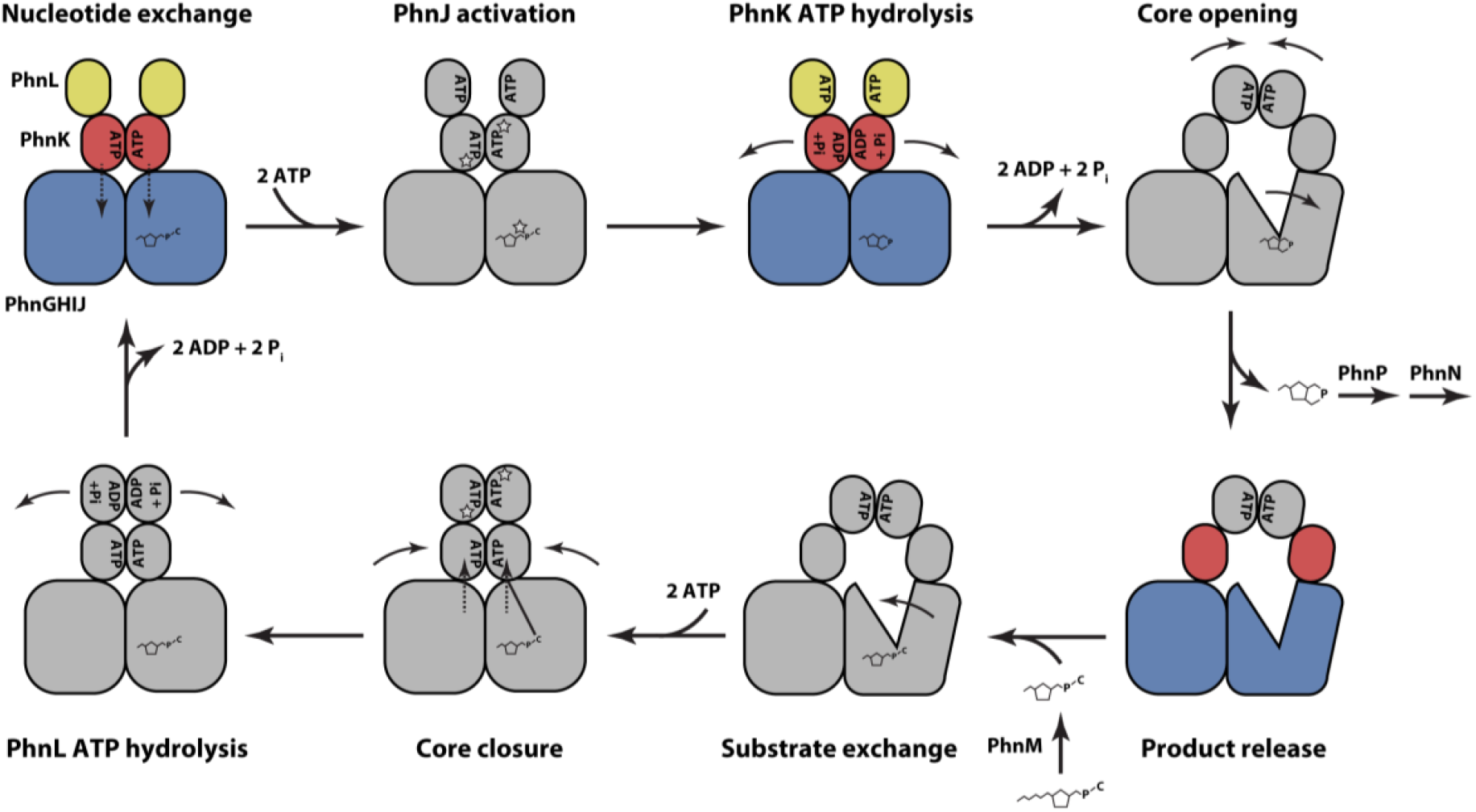
Model for the role of a dual ABC module in catalysis by C-P lyase. A schematic model depicting a possible catalytic cycle for C-P lyase with known (observed) states in colours and putative states in grey. At the top left, the C-P lyase core complex (PhnGHIJ, blue) binds two dimers of PhnK (red, in the ATP-bound state) and PhnL (open) and substrate in the form of a 5-phosphoribosyl-α-1-phosphonate. This complex might bind ATP in PhnL as well before triggering C-P bond cleavage in PhnJ leading to production of 5-phosphoribosyl-1, 2-cyclic phosphate. Next, following ATP hydrolysis and P_i_ release in the PhnK dimer, the core opens to allow product release while the PhnL subunits form a closed dimer, and exchange with a new 5-phosphoribosyl-α-1-phosphonate substrate generated by PhnM by cleavage of the pyrophosphate from a similar triphosphate. Following substrate binding, the core closes again, bringing the two PhnK subunits in close proximity in an ATP bound state. In the same step or following this, PhnL hydrolyses ATP, releasing ADP + P_i_ and separates leading back to the starting position.

The reaction mechanism of the bacterial C-P lyase machinery has remained an unresolved conundrum in biology since its discovery more than 35 years ago.^5, 16^ The large complexity of the pathway consisting of many interacting protein domains and enzymes, their internal dependencies, and requirement for anaerobic reaction conditions has proved a challenging combination. However, steady progress has been made in recent years, both from the biochemical and structural side, and we are now slowly beginning to reveal some of the inner secrets of this fascinating molecular system.^3, 4, 11^ The structural rearrangements of the C-P lyase core complex we present here are intriguing and the fact that there are two ABC engines connected but only one catalyses ATP hydrolysis *in vitro* suggests that more such unique states are yet to be revealed. Moreover, it is possible that the functioning of C-P lyase is connected to its intracellular localisation. Therefore, to uncover even more secrets of this important pathway, we need to go as close as possible to the natural state and study the reaction as it takes place inside cells.

## Acknowledgements

This work benefited from access to the Netherlands Centre for Electron Nanoscopy (NeCEN) at Leiden University, an Instruct-ERIC centre with assistance from Ludo Renault, the Electron Bio-Imaging Centre (eBIC) at Diamond Light Source, UK, and at the Danish National Cryo-EM Facility (EMBION) Aarhus node (iNANO, Aarhus University, Denmark). The authors are also thankful to the Aarhus University Electron Microscopy Computing Cluster (EMCC) and Thibaud Louis Antoine Dieudonné for help with the ATPase assay. Financial support was provided by Instruct-ERIC (PID 1514) and a grant from the Novo Nordisk Foundation (grant no. NNF18OC0030646) to D.E.B.

## Author Contributions

S.K.A, N.S., B.H.J., and D.E.B. designed and S.K.A, N.S, J.J.E. and R.B.S. carried out the experiments. N.S., S.K.A., and T.B. collected EM data, N.S. determined initial EM structures of Phn(GHIJ)_2_K to lower resolution with help from J.L.K. while S.K.A. determined all high-resolution structures and carried out structure refinement and validation. S.K.A. and D.E.B. wrote the manuscript draft and produced figures, and all authors revised the text.

## Additional Information

Atomic coordinates have been deposited in the Protein Data Bank (PDB) with accession codes 7Z19 (Phn(GHIJ)_2_K), 7Z16 (Phn(GHIJKL)_2_ PhnK-E171Q:AMPPNP), 7Z15 (Phn(GHIJKL)_2_ WT:ADP + P_i_), 7Z18 (Phn(GHIJK)_2_ WT:ATP closed), and 7Z17 (Phn(GHIJK)_2_ WT:ATP open). EM density maps have been deposited in the Electron Microscopy Data Bank (EMDB) with accession codes EMDB-14445 (Phn(GHIJ)_2_K), EMDB-14442 (Phn(GHIJKL)_2_ PhnK-E171Q:AMPPNP), EMDB-14441 (Phn(GHIJKL)_2_ WT:ADP + P_i_), EMDB-14444 (Phn(GHIJK)_2_ WT:ATP closed), and EMDB-14443 (Phn(GHIJK)_2_ WT:ATP open). The authors declare no competing financial interests. Correspondence and requests for materials should be addressed to D.E.B.

## Methods

### Expression and purification of Phn(GHIJ)_2_K

Plasmid pHO575 encoding PhnGHJIK with a C-terminal His-tag on PhnK was introduced into *E. coli* strain HO2735 *(Δ(lac)X74 ΔphnCDEFGHIJKLMNOP* 33–30/F lacI^q^ zzf::Tn10) as described previously^4^. The cells were grown in Luria Bertani (LB) media containing ampicillin (100 μg/mL) in a shaking incubator at 37°C until an OD_600_ of 0.5 was reached at which time gene expression was induced using 0.5 mM Isopropyl β-D-1-thiogalactopyranoside (IPTG). The cells were incubated overnight at 20°C, collected by centrifugation, resuspended in 50 mM HEPES/NaOH pH 7.5, 150 mM NaCl, 20 mM Imidazole, 5 mM MgCl_2_, 3 mM BME, 20% (v/v) glycerol, and 1 mM PMSF, lysed by sonication, and centrifuged at 23.400x g for 45 min at 4°C. The supernatant was loaded onto a 5 mL pre-equilibrated His-Trap HP column (Cytiva). The column was washed with 50 mM HEPES/NaOH pH 7.5, 650 mM NaCl, 20 mM imidazole, 5 mM MgCl_2_, 3 mM BME, and 20% (v/v) glycerol followed by 50 mM HEPES/NaOH pH 7.5, 150 mM NaCl, 20 mM imidazole, 5 mM MgCl_2_, 3 mM BME, 20% (v/v) glycerol, and 50 mM imidazole before elution in the same buffer with 300 mM imidazole. Purified samples were loaded onto a 1 mL Source 15Q (Cytiva) pre-equilibrated with 50 mM HEPES/NaOH pH 7.5, 5 mM MgCl_2_, 5 mM BME, and 100 mM NaCl, washed and eluted using a linear gradient from 100-800 mM NaCl over 20 column volumes (CVs). Samples containing PhnGHIJK were diluted to ~100 mM NaCl before loading onto a MonoQ column, pre-equilibrated in 50 mM HEPES/NaOH pH 7.5, 5 mM MgCl_2_, 5 mM BME, and 100 mM NaCl. Protein was eluted using a gradient from 100-500 mM NaCl over 36 CVs. Individual peaks were pooled and concentrated using Vivaspin 20 Ultrafiltration unit and loaded on a Superdex 200 Increase 10/300 GL size exclusion column preequilibrated in 50 mM HEPES pH 7.5, 125 mM NaCl, and 5 mM BME. Samples were stored on ice until used for cryo-EM grids.

### Cryo-EM data collection and analysis of Phn(GHIJ)_2_K

Cryo-EM grids (C-Flat 1.2/1.3 400 mesh copper grids, Protochips) were glow dischargied for 90 seconds at 10 mA using a GloQube system (Quorum Technologies) immediately before applying 3 μL of purified Phn(GHIJ)_2_K (1.7 mg/mL), blotting (6-7 s) and plunge-freezing in liquid ethane using a Leica plunge freezer EM GP2, set to 5°C, 100% humidity and an ethane temperature of −184°C. EM data were collected at the Electron Bio-Imaging Centre (eBIC) at Diamond Light Source, UK on a Titan Krios 300 keV with a direct electron detector (Gatan K3), an energy filter operated with a slit width of 20 eV and using SerialEM^27^ for data acquisition. 17,088 movies were collected with an exposure of 0.2 seconds per frame over 40 frames with a dose of 1.3 e^-^/Å^2^/frame corresponding to a total exposure of 52 e^-^/Å^2^, the defocus ranging from −0.7 to −2.0 μm the nominal magnification was set to 135.000 giving a physical pixel size of 0.83 Å in counting mode. Data collection quality was monitored by running RELION^28^ in streaming mode through relion_it.py.

Frame-based motion correction was performed using RELIONs implementation of MotionCor2^29^ and CTF estimation were performed using Gctf^30^ within RELION-3. Subsets of the micrographs were used for Laplacian-of-Gaussian picking in RELION-3 before extraction with 8.75 times binning and 2D classification. Good 2D classes were used as templates template picking. Picked particles were extracted using 6 times binning and 2D classification to remove bad particles. Particles were reextracted using 4 times binning, followed by initial model generation and 3D classification. The resulting 3D classes resembling PhnGHIJK contained a stack of 1,329,400 particles. A second round of 3D classification in RELION-3 and heterogenous refinement in cryoSPARC (Structura Biotechnology Inc.)^31^ led to a stack of 901,800 particles. These particles were moved back into RELION-3.1 where they were further refined using Bayesian polishing, per-particle CTF refinement, estimation of higher-order aberrations, beam tilt and anisotropic magnification^32^. After autorefinement in RELION-3.1, an FSC resolution with masking of 2.18 Å was reached. The resulting map showed great detail of the PhnGHIJ part of the structure which agreed well with the former published structure^4^, though the PhnK part of the structure had low local resolution. To improve the interpretation of PhnK, 3D classification with signal subtraction and 3D variability was used^19, 20^ as detailed in **Supplementary Figure 2**. Particles from resulting classes showing interpretable density for PhnK were reextracted and used for new 3D auto-refinement runs. The resulting map had a FSC resolution of 2.57 Å and improved local resolution for PhnK. An initial model was constructed by using the published structure of Phn(GHIJ)_2_ (PDB: 4XB6)^4^ and a predicted model of PhnK from Phyre2^33^ for rigid-body modelling using UCSF Chimera.^34^ This model was further fitted into the map using Namdinator^35^ and manually built in Coot^36^ for the regions of the map with higher resolution whereas ISOLDE^37^ was used for regions of lower resolution and interpretability. The final model was refined and validated using Phenix real-space refinement^38^.

### Cloning, expression and purification of Phn(GHIJKL)_2_ PhnK E171Q

pRBS01 was constructed using Gibson assembly^39^ by extraction of *phnGHIJKLMNOP* from pBW120^40^ and insertion of a sequence encoding a TEV-2xStrep-tag at the 3’ end of *phnK.* Linearized pET28(a) vector was prepared by PCR, using primers RBS01 and RBS02, while the TEV-2x-Strep site was prepared from a gBlock sequence using primers RBS03 and RBS04. The two fragments containing the *phn* genes were prepared from pBW120 using primers RBS05 + RBS06 *(phnGHIJK)* and RBS07 + RBS08 *(phnLMNOP*), respectively. Linearised plasmid was mixed with 2-3-fold excess of the three insert fragments followed by addition of Gibson Assembly® Master Mix (New England Biolabs) and incubation at 50°C for 60 minutes. Assembled plasmids were transformed into *E. coli* strain NEB 10-β competent cells (New England Biolabs) and plated on LB agar containing 50 μg/mL kanamycin. The PhnK E171Q mutant was constructed by PCR using primers SKA01 and SKA02. pRBS01 PhnK E171Q was transformed into competent *E. coli* Lemo21 cells (New England Biolabs). Cells were grown in LB media containing 300 μM L-rhamnose, chloramphenicol (34 μg/mL) and kanamycin (50 μg/mL) to an OD_600_ of ~0.5 before addition of 1 mM IPTG. The cells were incubated at 37°C for 3-4 h before harvesting. Collected cells were resuspended in lysis buffer containing 25 mM HEPES/KOH pH 7.5, 125 mM KCl, 5 mM MgCl_2_, 10% glycerol, 5 mM BME, 1 μg/mL DNase I, 1 mM PMSF, and 5 μM adenosine-5’-[(β-γ)-imido]triphosphate (AMPPNP) and lysed by sonication. The lysate was cleared by centrifugation at 23.400x g for 45 min at 4°C and loaded onto a 1 mL Strep-Trap HP column (Cytiva) which were kept at 8°C and had been pre-equilibrated in lysis buffer. The column was washed with 10 CV of a Strep-Trap wash buffer containing 25 mM HEPES/KOH pH 7.5, 125 mM KCl, 5 mM MgCl_2_, 5 mM BME, and 5 mM AMPPNP before elution with 5 CV of this buffer plus 2.5 mM *D*-desthiobiotin. All fractions were analysed by SDS-PAGE and stored on ice.

### Cryo-EM sample preparation and analysis of Phn(GHIJKL)_2_ PhnK E171Q

UltrAuFoil 0.6/1 grids were glow discharged (10 mA, 90 sec, GloQube, Quorum) immediately before dispensing 3 μL freshly purified complex diluted in Strep-Trap wash buffer to a concentration of 2.5 mg/mL followed by blotting (7-9 s) and immediate plunge-freezing in liquid ethane using a Leica plunge freezer EM GP2, adjusted to 5°C, 100% humidity and an ethane temperature of −184°C. Cryo-EM data was acquired at the Danish National Cryo-EM Facility (EMBION) – AU node (iNANO, Aarhus University) on a Titan Krios G3i (Thermo Fisher Scientific) operated at 300 keV using a K3 direct electron detector (Gatan), a Bioquantum energy filter (Gatan) with a slit width of 20 eV and EPU (Thermo Fisher Scientific) for automated data collection. Data were collected in counted super resolution mode at a nominal magnification of 130,000 with gain correction on the fly followed by 2x binning in EPU resulting in movies output at physical pixel size (0.647 Å/pixel). Eucentric height was determined for each grid square and data acquired using a defocus range of −0.5 to −1.4 μm and an electron fluence of 62 e^-^/Å^2^ over 56 frames. Data quality was monitored and initially processed through cryoSPARC Live.

All data were processed inside cryoSPARC. Initially, micrographs were patch motion corrected, followed by patch CTF-estimation and filtering using various data quality indicators. Particles were initially picked using a gaussian blob picker, after which 4x binned particles were extracted and subjected to 2D classification. Particles from good 2D classes were kept and a manual picking job using a subset of 14 micrographs was initiated where the remaining particles were picked. The picked particles were used to train the Deep picker, after which the model was used to pick particles the remaining 4,502 micrographs.

The data processing strategy is shown in **Supplementary Figure 4**. Briefly, a total of 1,530,855 particles were extracted using 4x binning and subjected to 2D classification, following selection of good 2D classes and 3D *ab initio* reconstruction using five classes, followed by heterogenous refinement using the same classes. A class showing the PhnGHIJKL structure was subjected to a homogenous refinement and the aligned particles was used for 3D variability analysis.^20^ For 3D variability analysis, three orthogonal principal modes were solved with resolution filtered to 5.5 Å, particles from different conformations were separated by clustering the particles using their principal modes into four different clusters. For the PhnGHIJKL PhnK E171Q structure, one cluster was selected and particles from this class was used for local motion correction following extraction in a 1.48x binned 420×420 pixel box, which were used for refinement with C2-symmetry imposed giving a final FSC resolution of 2.09 Å resolution from 59,737 particles. This map was further processed by determining the local resolution and during local filtering and sharpening of the map using the local resolution data. For model building, the PhnGHIJ core complex and PhnK from the Phn(GHIJ)_2_K structure were individually docked into the map using ChimeraX^41^ while a PhnL model was generated using Phyre2.^33^ The model was further improved by map fitting using Namdinator^35^, following correction of the individual chains and addition of ligands and ions using ISOLDE^37^. The model was inspected in Coot^36^ where additional residues were added when visible in the map. The final model was refined and validated using Phenix real-space refinement^42^.

### Purification of wild type Phn(GHIJKL)_2_

Expression and purification of Phn(GHIJKL)_2_ with wild type PhnK from pRBS01 followed the same protocol as the PhnK E171Q mutant. Post StrepTrap purification, the sample was treated with TEV protease (1:50 w/w TEV protease) at 8°C overnight, after which the sample was loaded on a MonoQ 5/50 column (Cytiva), pre-equilibrated in 25 mM HEPES/KOH, 100 mM KCl, 5 mM MgCl_2_, 10% glycerol, 5 mM BME, and 1 μM AMPPNP). The protein was eluted using a gradient of 200-450 mM KCl over 36 CVs after which individual peaks were pooled. The Phn(GHIJKL)_2_ sample was concentrated using Vivaspin 20 Ultrafiltration unit and loaded on a Superdex 200 increase 10/300 column (Cytiva) equilibrated in 25 mM HEPES/KOH, 125 mM KCl, 0.5 mM EDTA, 5 mM BME, and 0.25 mM ATP. Purified protein was concentrated and stored at 4°C.

### ATPase assays

Purified Phn(GHIJKL)_2_ complexes were incubated at a concentration of 210 nM with 5 mM ATP in 50 mM HEPES/KOH pH 7.5, 300 mM KCl, 10 mM MgCl_2_, and 5 mM BME at 30°C overnight. 100 μL of a 1:20 dilution of the reaction mixture was loaded onto a MonoQ column (Cytiva) pre-equilibrated in 25 mM HEPES/KOH pH 7.5, 5 mM MgCl_2_, and 5 mM BME and eluted using a gradient from 0-180 mM KCl over 7 CVs. The column was furthermore calibrated with known samples of ATP, ADP, AMP, and adenine to allow identification of potential reaction products. The quantitative rate of ATPase hydrolysis was measured via a coupled enzyme assay conducted at 37°C in 25 mM HEPES/KOH, 125 mM KCl, 0.5 mM EDTA, 5 mM BME, and 0.25 mM ATP with the addition of 1 mM phosphoenolpyruvate, 350 μM NADH, 0.04 mg/mL pyruvate kinase, 0.1 mg/mL lactate dehydrogenase, 10 mM MgCl_2_, and ATP to a final concentration of 1 or 5 mM. The reaction was initiated by addition of 0.02 mg purified wild type or PhnK E171Q mutant complex in a total volume of 0.35 mL resulting in a final protein concentration of 0.057 mg/mL. Conversion of ATP to ADP was calculated from the measured NADH oxidation rates using an extinction coefficient of 6.2 L*mM^-1^*cm^-1^ at 340 nm. The reaction was followed for 6 minutes, and the reaction rate was measured using a linear fit to the degradation of NADH to NAD^+^.

### Cryo-EM sample preparation and analysis of wild type Phn(GHIJKL)_2_ under ATP turnover conditions

A sample of purified Phn(GHIJKL)_2_ (2.5 mg/mL) was kept on ice and incubated with ATP at a concentration of 1.5 mM followed by addition of MgCl_2_ to a concentration of 6 mM to activate the ATPase reaction. Samples were incubated for 15 s at 37°C and immediately plunge-frozen onto glow discharged (10 mA, 90 sec, GloQube, Quorum) UltrAuFoil 0.6/1 grids using a Leica plunge freezer EM GP2 set to 37°C, 100% humidity and an ethane temperature of −184°C and a blotting time of 6-9 seconds. Data were acquired in a similar way as for PhnK E171Q. All data were processed inside cryoSPARC. Initially micrographs were patch motion corrected, followed by patch CTF-estimation and filtered using various data quality indicators. Particles were initially picked using a gaussian blob picker, after which 4x binned particles were extracted and subjected to 2D classification. Particles from clear 2D classes were kept and a manual picking job using a subset of 14 micrographs was initiated where the remaining particles were picked. The picked particles were used to train the Deep picker, after which the model was used to pick particles the remaining 3,741 micrographs.

Initial data processing was conducted as for the Phn(GHIJKL)_2_ Phn E171Q dataset, leading to 1,155,385 particles picked form 3,741 micrographs. Data processing strategy is shown in **Supplementary Figure 8 and 9**. Picked particles were extracted using 2.75x binning and subjected to 2D classification, following selection of good 2D classes. The selected particles from 2D classification were used for 3D *ab initio* reconstruction using four classes. Particles from classes showing protein-like features were then used for non-uniform refinement.^43^ The resulting particles from the non-uniform refinement were subjected to 3D variability analysis^20^ to separate different conformations from each other using three orthogonal principal modes that were solved with resolution filtered to 6 Å, after which particles from different conformations were separated by clustering according to the obtained principal modes, into 8 different clusters. For the Phn(GHIJKL)_2_ WT structure, two clusters were selected, and the corresponding particles were subjected to perparticle local motion correction and reextracted in a 1.29x binned 480×480 pixel box. The final set of 222,056 particles was then used for non-uniform refinement using C2 symmetry to gain the final map with a FSC resolution of 1.93 Å (**Supplementary Figure 9, top**).

For the Phn(GHIJK)_2_ open conformation, four clusters from the first round of 3D variability were subjected to another round of 3D variability and clustering. One cluster was selected, and the particles were reextracted in a 1.42x binned 386×386 pixel box. The final set of 31,280 particles was used for non-uniform refinement and gave a map with a FSC resolution of 2.57 Å (**Supplementary Figure 9, bottom**). This map was further processed by determining the local resolution and during local filtering and sharpening of the map using the local resolution data. For the Phn(GHIJK)_2_ closed conformation one cluster was selected and the particles was reextracted in a 1.42x binned 386×386 pixel box. The final set of 81.605 particles were used for non-uniform refinement with C2 symmetry and gave a map with a FSC resolution of 1.98 Å (**Supplementary Figure 9, middle**).

For model building of Phn(GHIJKL)_2_ WT, the final Phn(GHIJKL)_2_ PhnK E171Q model was initially docked into the map. Ligands were exchanged, waters were added using Phenix Douse^38^, and the model built using Coot and ISOLDE. The final model was refined and validated using Phenix real-space refinement. The Zn^2+^ binding site at His 328 and 333 in PhnI, ligand bond lengths was restrained using data from C. Lim and T. Dudev, 2000^44^. For the Phn(GHIJK)_2_ open conformation, the initial model was based on the wild type Phn(GHIJKL)_2_ structure, which was split into two parts, one containing PhnG2HI2JK and another containing PhnHJK, each representing one side of the open core complex, which were docked into the map using ChimeraX. The model was built using ISOLDE and Coot, except for the two PhnK molecules and part of one of the PhnJ molecule, which were not modified due to low local resolution. The final model was refined and validated using Phenix real-space refinement, without refinement of the PhnK chains. For the Phn(GHIJK)_2_ closed conformation the model of Phn(GHIJKL)_2_ structure without PhnL was docked into the map and ligands were exchanged and waters removed. The final model was refined using Phenix real-space refinement.

### Complementation of *Δphn* by *phnJ, phnK* and *phnL* mutation

For complementation, a plasmid (pSKA03) containing the entire *phn* operon including the *pho* box was constructed based on pBW120 by removal of non-essential sequences. Initially, two large fragments were created using PCR, a plasmid backbone with primers SKA03 and SKA04 and another fragment containing the whole *phn* operon including the *pho* box using primers SKA05 and SKA06. The products were treated with restriction endonuclease DpnI and transformed together (300 ng each) into competent *E. coli* NovaBlue cells to achieve RecA-independent cloning^45^. Plasmid DNA was purified from positive clones and confirmed by nucleotide sequencing. Subsequently, the PhnJ E149A, PhnJ Y158A, PhnK R78A/R82A, PhnK E171Q, and PhnL E175Q point mutations were introduced by PCR using primers SKA07 and SKA08 for PhnJ E149A, SKA09 and SKA10 for PhnJ Y158A, SKA11 and SKA12 for PhnK R78A/R82A, SKA13 and SKA14 for PhnK E171Q, SKA15 and SKA16 for PhnL E175Q. For complementation, the phosphonate deficient *E. coli* strain HO1488 *(ΔphnHIJKLMNOP)* was used. Strain HO1488 was grown in LB containing 50 μg/mL kanamycin to an OD_600_ of ~0.3, concentrated by centrifugation, and resuspended in 200 μL ice cold TSB (10% w/v PEG 3350, 5% DMSO, and 20 mM MgCl_2_ in LB media) to make the cells competent for DNA transformation by heat shock with DNA of the plasmids mentioned above. Transformed cells were grown on MOPS minimal agar plates^26^ containing 0.2% glucose, 100 μg/mL ampicillin, 34 μg/mL kanamycin, and 0.2 mM of either K_2_HPO_4_, 2-aminoethyl phosphonate, methyl phosphonate or no added phosphate source.

