## Supplemental Data for "Structural remodelling of the carbon-phosphorus lyase machinery by a dual ABC ATPase"

SUPPLEMENTARY DATA

**Supplementary Figure 1**

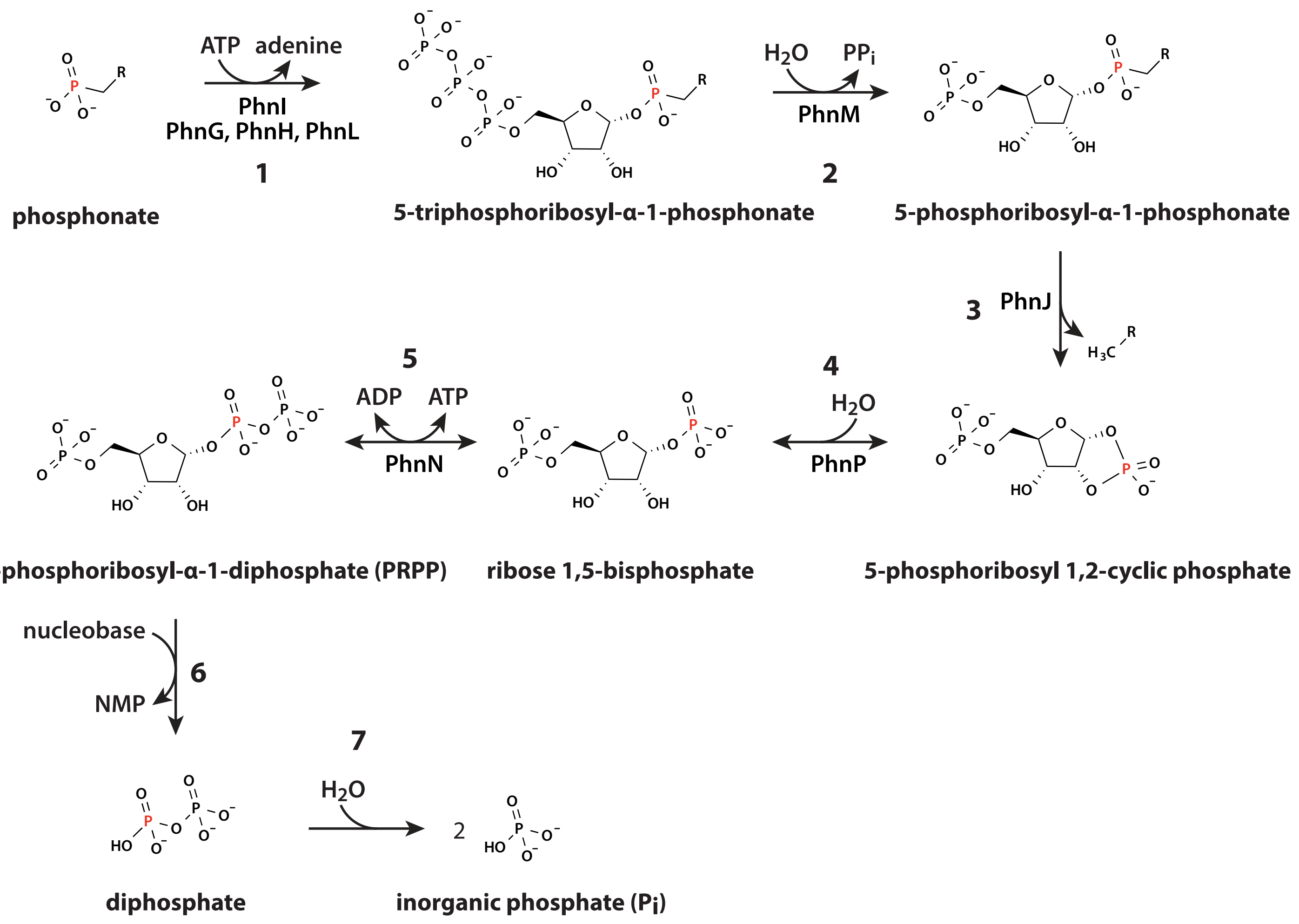

**Supplementary Figure 2**

**Dataset I: Phn(GHIJ)$_2$K wild type**

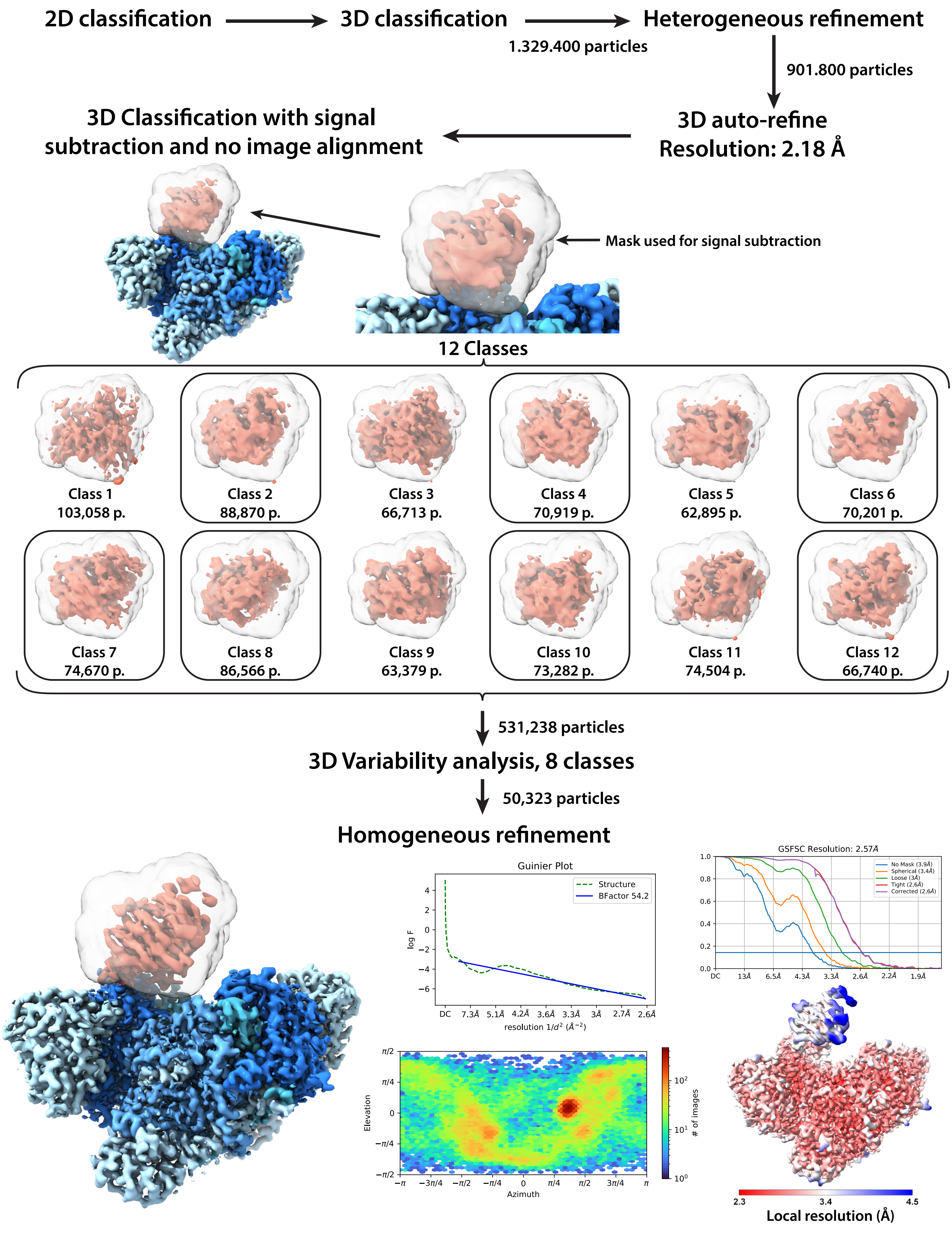

### Supplementary Figure 3

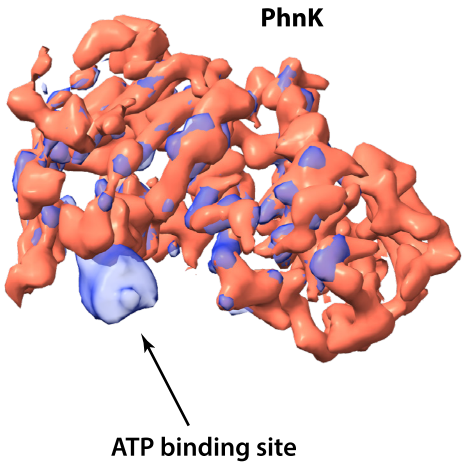

**Supplementary Figure 4**

**Dataset II: Phn(GHIJKL)$_2$ PhnK-E171Q + AMPPNP**

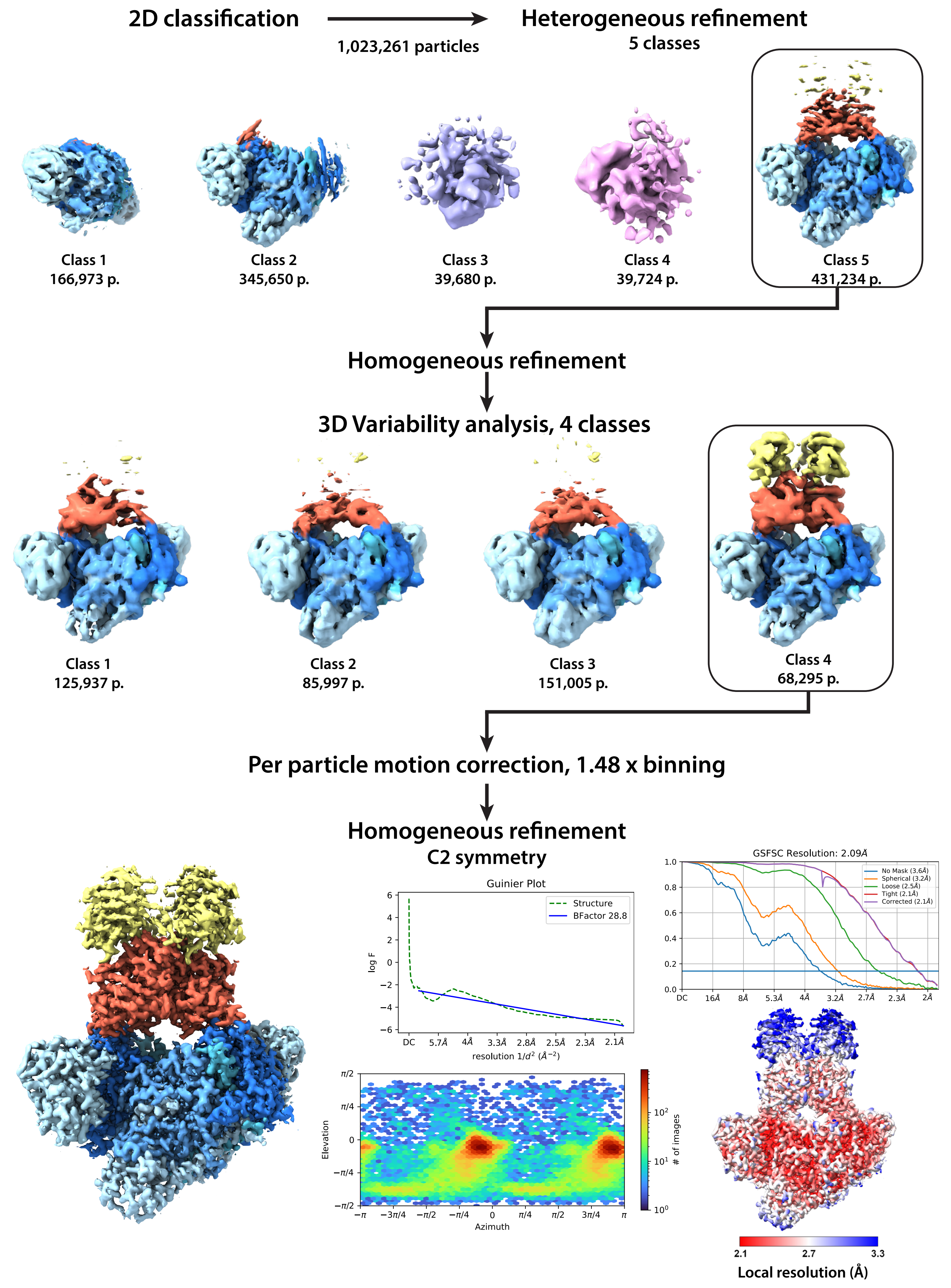

#### Supplementary Figure 5

**A**

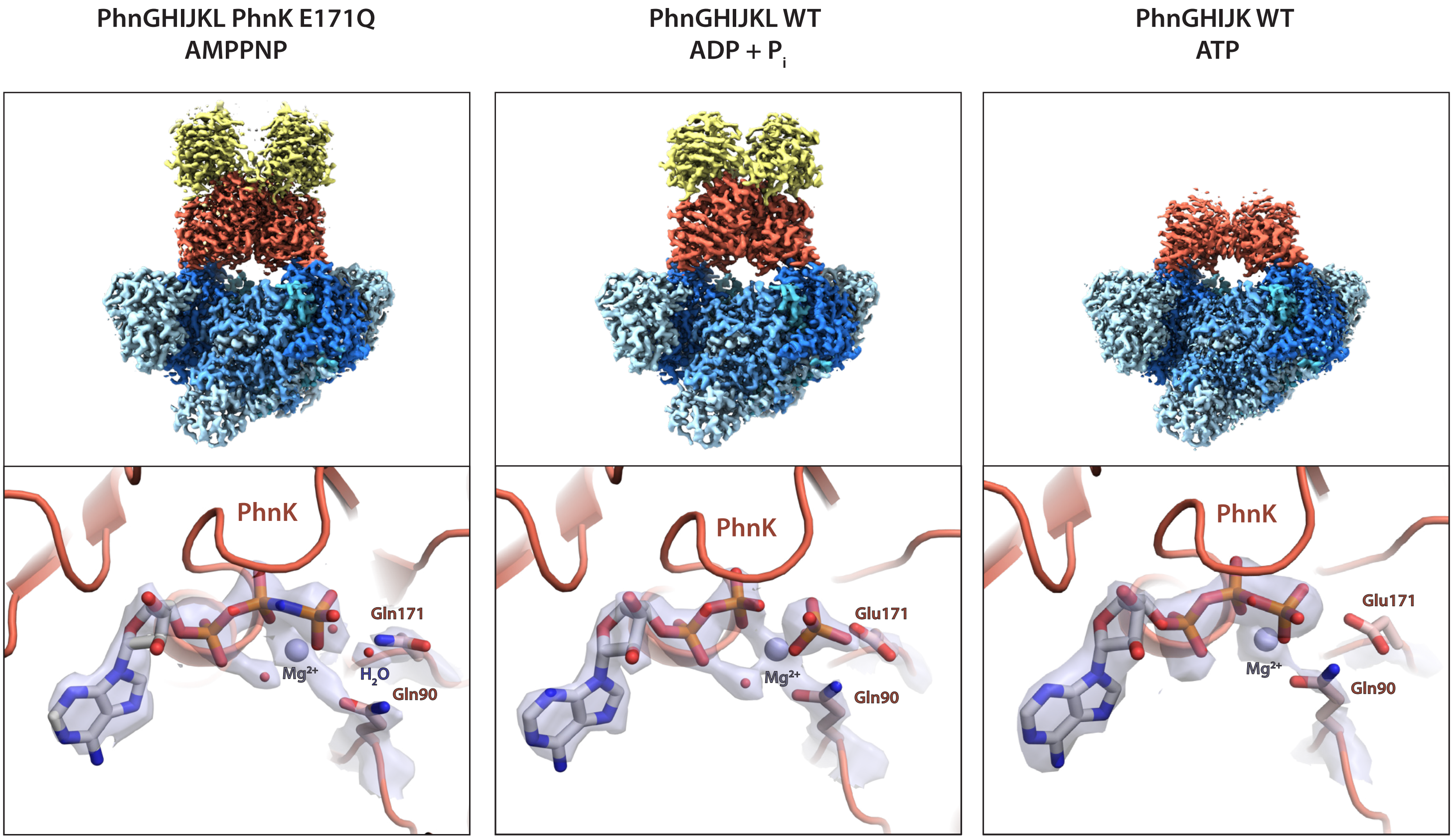

**B**

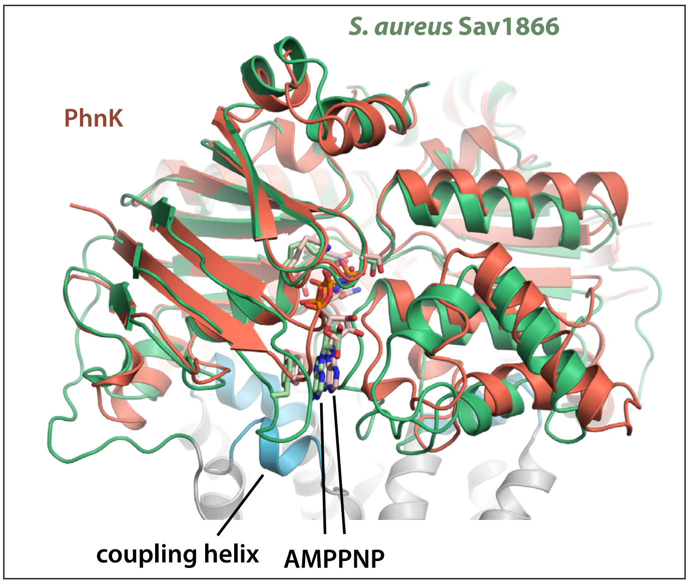

**C**

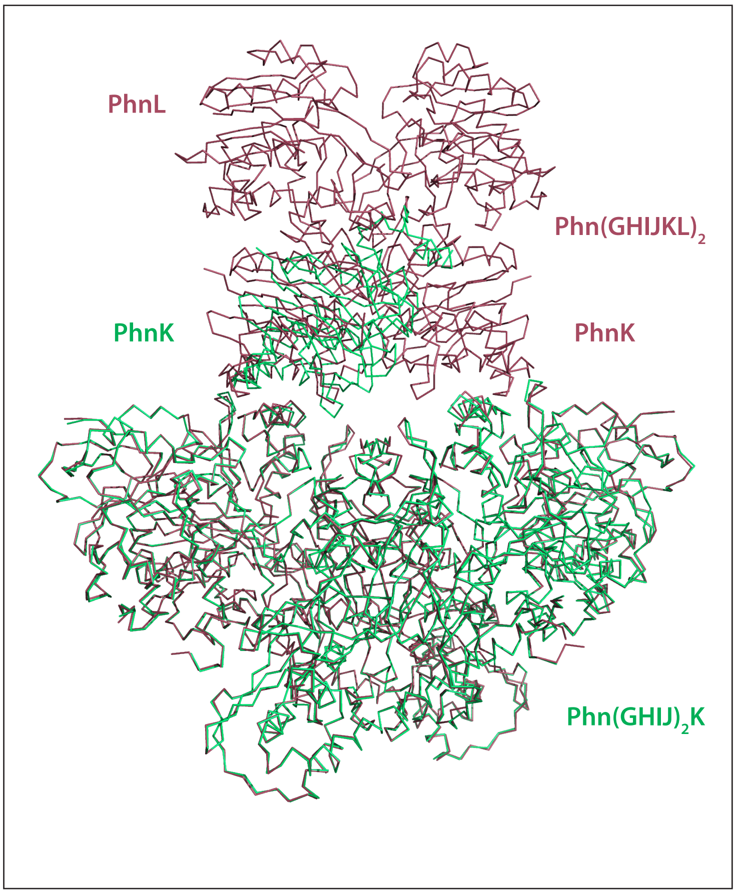

### Supplementary Figure 6

## A

| emPAI | Cov | ↑*Family* | M | DB | Accession | Score | Mass | Matches | Match(sig) | Sequences | Seq(sig) | Description |
|---|---|---|---|---|---|---|---|---|---|---|---|---|
| 16.41 | 0.57 | 1 | 1 | Trembl | **A0A023Z6E0_ECOLX** | 13332 | 32067 | 375 | 375 | 14 | 14 | Alpha-D-ribose 1-methylphosphonate 5-phosphate C-P lyase OS=Escherichia coli O145:H28 str. RM12581 OX=1248823 GN=**phnJ** PE=3 SV=1 |
| 13.98 | 0.57 | 1 | 2 | Trembl | **A0A6G6LIX9_ECOLX** | 13200 | 32067 | 367 | 367 | 13 | 13 | Alpha-D-ribose 1-methylphosphonate 5-phosphate C-P lyase OS=Escherichia coli OX=562 GN=**phnJ** PE=3 SV=1 |
| 6.83 | 0.37 | 1 | 3 | Trembl | **A0A2T3V5M7_ECOLX** | 11531 | 47038 | 348 | 348 | 15 | 15 | ATP-binding cassette domain-containing protein OS=Escherichia coli OX=562 GN=C9E25_06215 (**phnK**) PE=4 SV=1 |
| 3.50 | 0.40 | 1 | 4 | Trembl | **A0A376IAT9_ECOLX** | 9646 | 32095 | 273 | 273 | 8 | 8 | Alpha-D-ribose 1-methylphosphonate 5-phosphate C-P lyase OS=Escherichia coli OX=562 GN=**phnJ** PE=3 SV=1 |
| 13.60 | 0.34 | 1 | 5 | Trembl | **A0A6N0FP14_ECOLX** | 7645 | 37827 | 268 | 268 | 13 | 13 | Alpha-D-ribose 1-methylphosphonate 5-triphosphate synthase subunit PhnI OS=Escherichia coli OX=562 GN=**phnI** PE=4 SV=1 |
| 6.96 | 0.42 | 1 | 6 | Trembl | **A0A7B1JLD3_ECOLX** | 4130 | 27813 | 133 | 133 | 9 | 9 | ABC transporter ATP-binding protein OS=Escherichia coli OX=562 GN=**phnK** PE=4 SV=1 |
| 3.87 | 0.31 | 1 | 7 | Trembl | **A0A788LHF4_ECOLX** | 3507 | 21197 | 116 | 116 | 6 | 6 | Phosphonate C-P lyase system protein PhnH OS=Escherichia coli OX=562 GN=**phnH** PE=4 SV=1 |
| 2.60 | 0.28 | 1 | 8 | Trembl | **A0A376RQC2_ECOLX** | 3327 | 56753 | 113 | 113 | 11 | 11 | Phosphonates transport ATP-binding protein PhnL OS=Escherichia coli OX=562 GN=**phnL** PE=4 SV=1 |
| 2.88 | 0.26 | 1 | 9 | Trembl | **A0A7B7R490_ECOLX** | 2963 | 21228 | 97 | 97 | 5 | 5 | Phosphonate C-P lyase system protein PhnH OS=Escherichia coli OX=562 GN=**phnH** PE=4 SV=1 |
| 2.10 | 0.26 | 1 | 10 | Trembl | **A0A7B6UGW1_ECOLX** | 1242 | 21154 | 40 | 40 | 5 | 5 | Phosphonate C-P lyase system protein PhnH OS=Escherichia coli OX=562 GN=**phnH** PE=4 SV=1 |
| 4.58 | 0.45 | 1 | 11 | Trembl | **A0A5C9AAH9_ECOLX** | 1181 | 16686 | 31 | 31 | 5 | 5 | ATP-binding cassette domain-containing protein (Fragment) OS=Escherichia coli OX=562 GN=FWK02_34920 (**phnL**) PE=4 SV=1 |
| 3.15 | 0.29 | 1 | 12 | Trembl | **A0A418GIX9_ECOLX** | 1146 | 16803 | 38 | 38 | 4 | 4 | ATP-binding cassette domain-containing protein OS=Escherichia coli OX=562 GN=D3C88_16455 (**phnK**) PE=4 SV=1 |
| 63.30 | 0.23 | 1 | 13 | Trembl | **A0A7A1M1F4_ECOLX** | 648 | 8967 | 42 | 42 | 3 | 3 | Carbon-phosphorus lyase complex subunit **PhnI** (Fragment) OS=Escherichia coli OX=562 GN=HJ332_004423 PE=4 SV=1 |
| 55.86 | 0.61 | 2 | 1 | Trembl | **A0A376TPT2_ECOLX** | 4221 | 15267 | 160 | 160 | 9 | 9 | Phosphonate metabolism protein OS=Escherichia coli OX=562 GN=**phnG** PE=4 SV=1 |
| 12.18 | 0.43 | 2 | 2 | Trembl | **A0A449CRE9_ECOLX** | 3552 | 16642 | 133 | 133 | 7 | 7 | Carbon-phosphorus lyase complex subunit OS=Escherichia coli OX=562 GN=**phnG** PE=4 SV=1 |
| 0.66 | 0.13 | 3 | 1 | Trembl | **A0A023Z718_ECOLX** | 273 | 57464 | 7 | 7 | 6 | 6 | 60 kDa chaperonin OS=Escherichia coli O145:H28 str. RM12581 OX=1248823 GN=groL PE=3 SV=1 |
| 0.33 | 0.06 | 4 | 1 | Trembl | **A0A023Z3R1_ECOLX** | 64 | 16733 | 1 | 1 | 1 | 1 | Biotin carboxyl carrier protein of acetyl-CoA carboxylase OS=Escherichia coli O145:H28 str. RM12581 OX=1248823 GN=accB PE=4 SV=1 |
| 0.17 | 0.05 | 5 | 1 | Trembl | **A0A023YYV7_ECOLX** | 58 | 30545 | 1 | 1 | 1 | 1 | Protein MtfA OS=Escherichia coli O145:H28 str. RM12581 OX=1248823 GN=mtfA PE=3 SV=1 |
| 0.32 | 0.07 | 6 | 1 | Trembl | **A0A023Z5Q5_ECOLX** | 53 | 17464 | 1 | 1 | 1 | 1 | Regulator of ribonuclease activity A OS=Escherichia coli O145:H28 str. RM12581 OX=1248823 GN=menG PE=3 SV=1 |
| 0.63 | 0.09 | 7 | 1 | Trembl | **A0A0E0U6W5_ECOLX** | 49 | 9656 | 1 | 1 | 1 | 1 | Uncharacterized protein OS=Escherichia coli UMNK88 OX=696406 GN=UMNK88_5065 PE=4 SV=1 |
| 0.07 | 0.02 | 8 | 1 | Trembl | **A0A023YRN2_ECOLX** | 47 | 69130 | 1 | 1 | 1 | 1 | Chaperone protein DnaK OS=Escherichia coli O145:H28 str. RM12581 OX=1248823 GN=dnaK PE=2 SV=1 |

## B

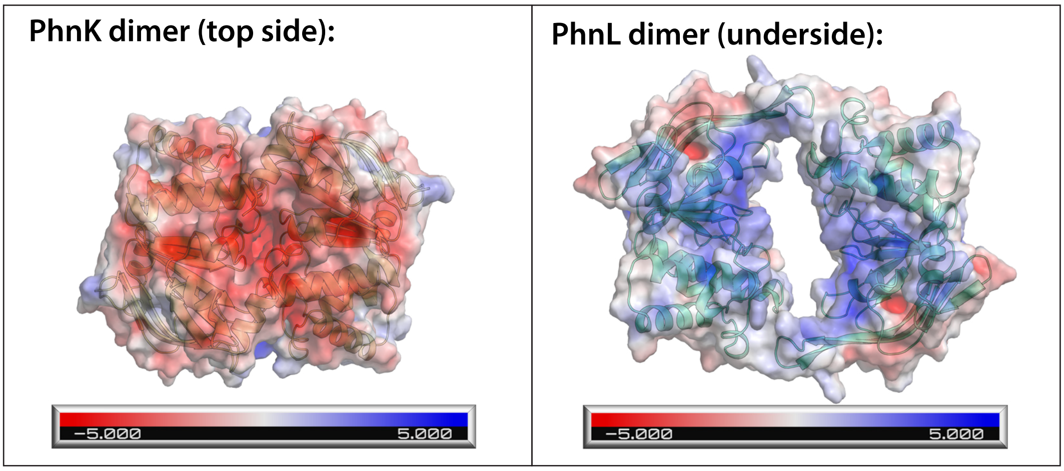

## C

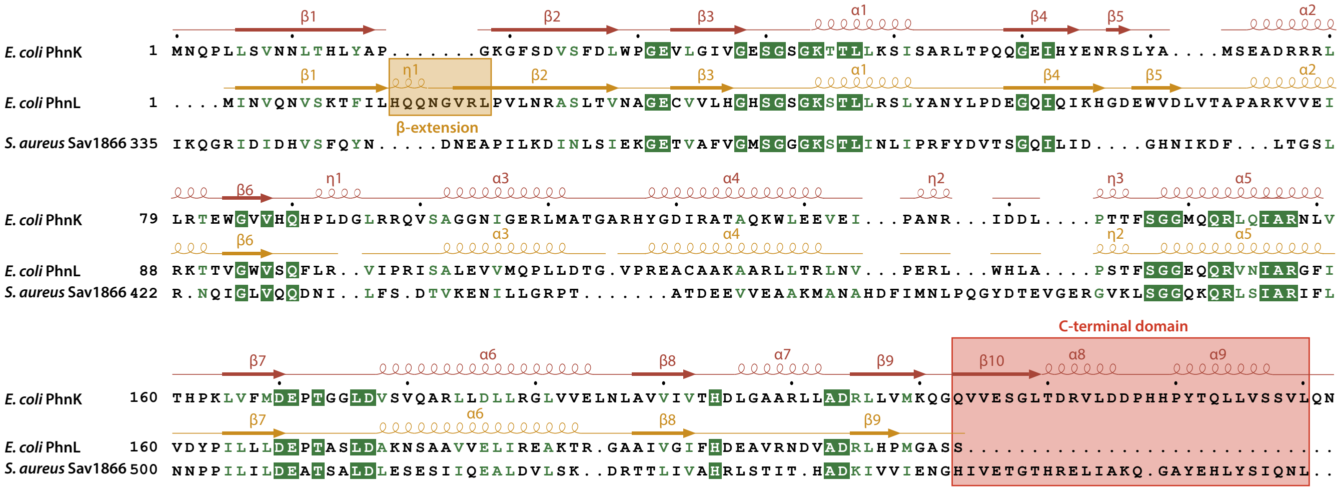

#### Supplementary Figure 7

**A**

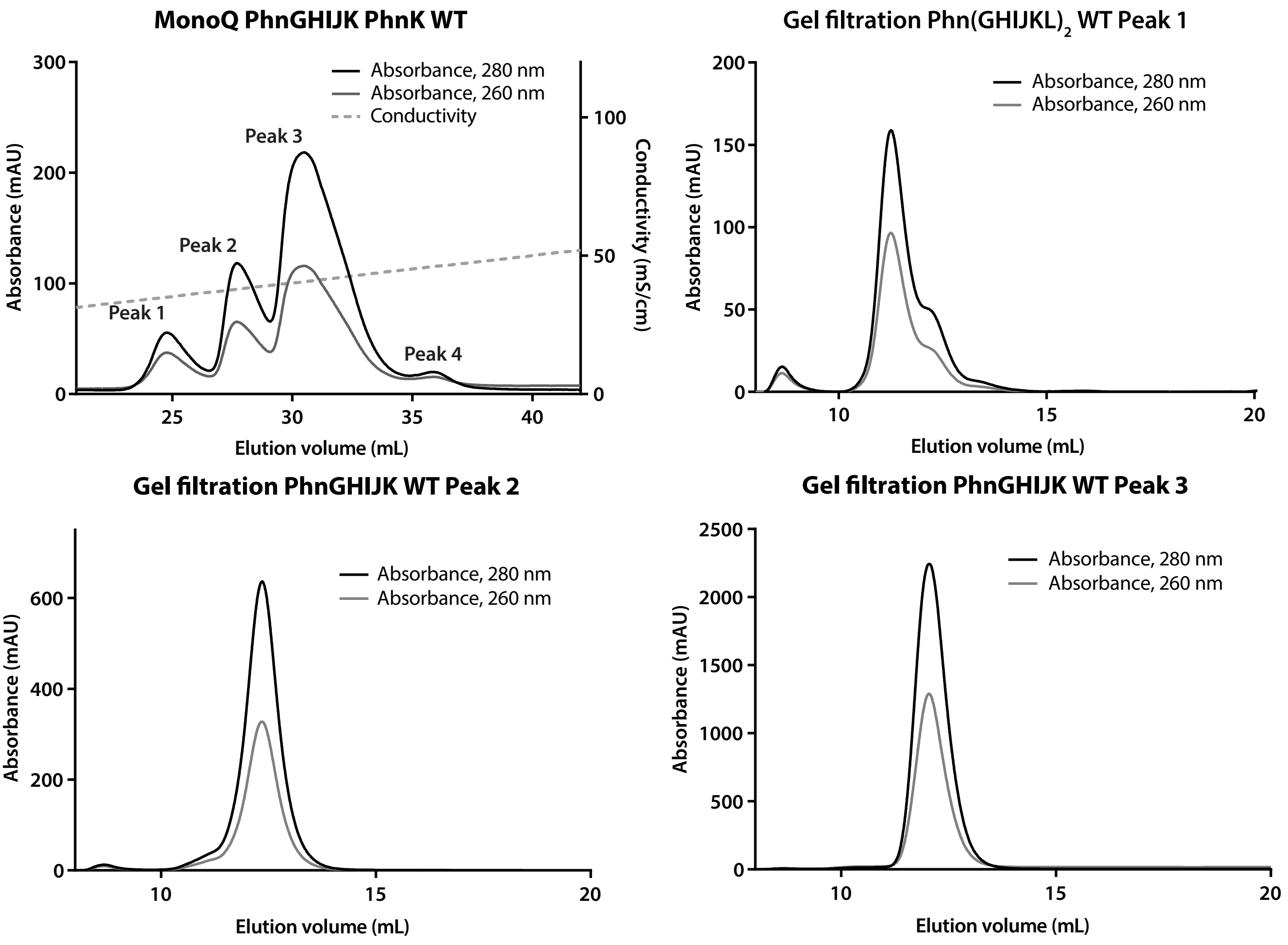

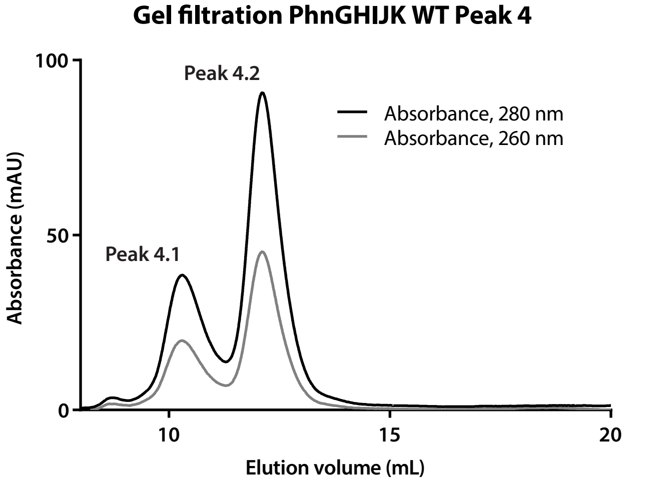

**B**

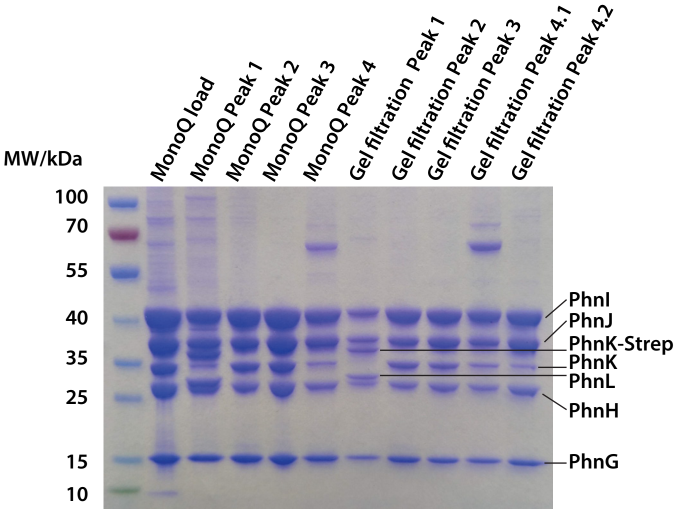

**Supplementary Figure 8**

Dataset III: Phn(GHIJKL)$_2$ WT under ATP turnover conditions

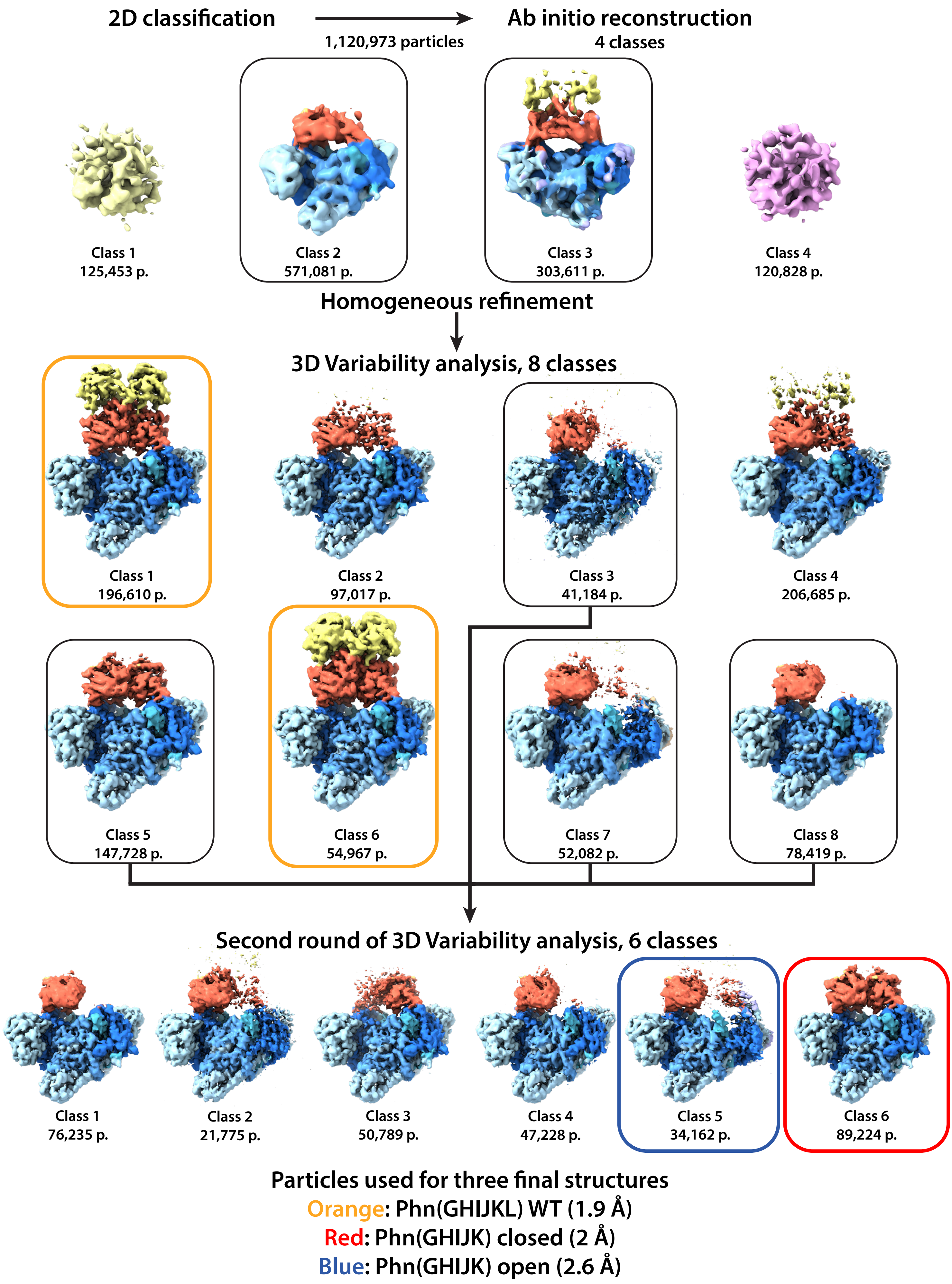

#### Supplementary Figure 9

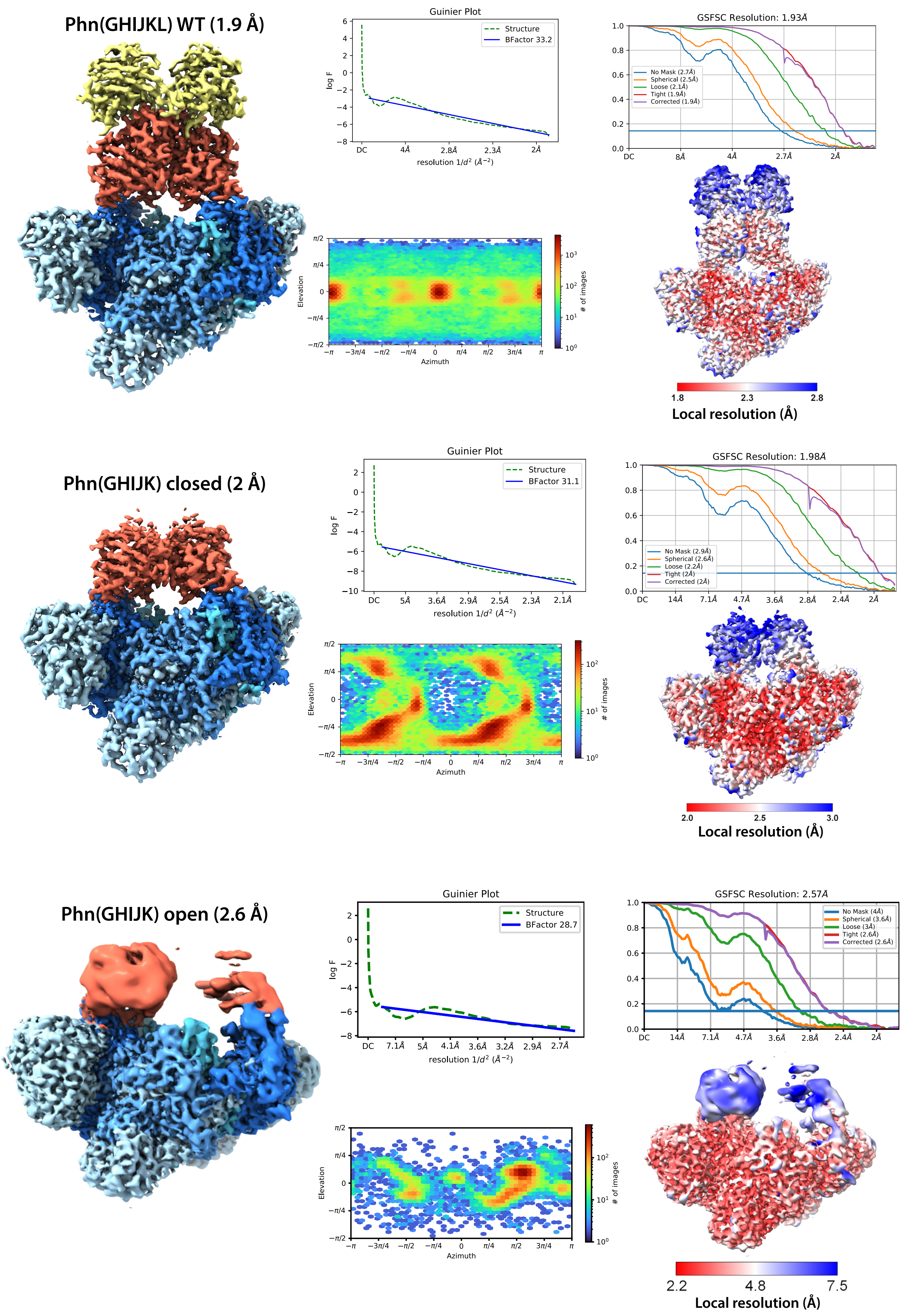

#### Supplementary Figure 10

A

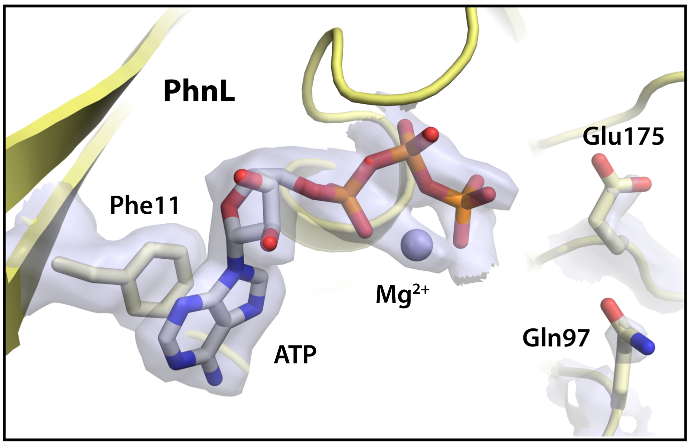

B

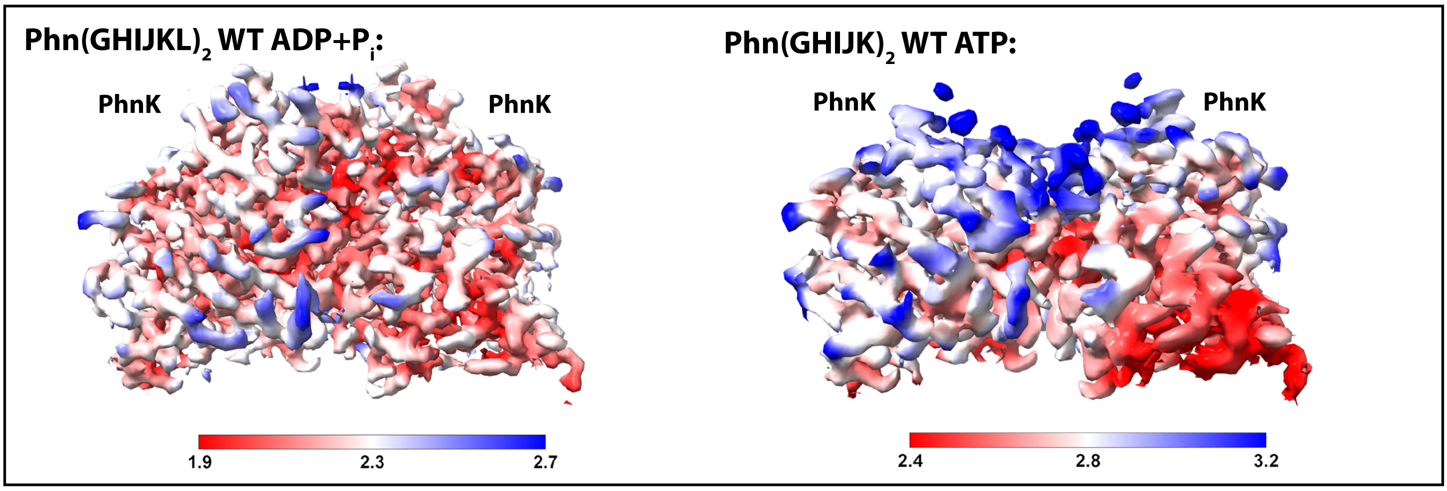

C

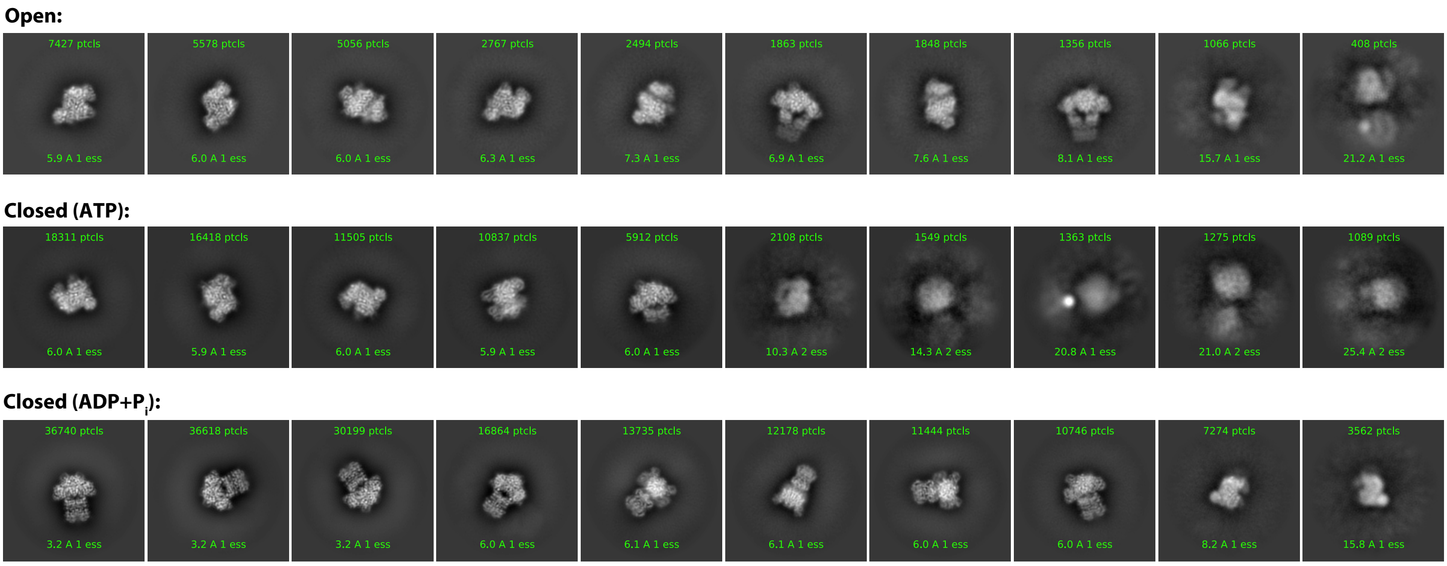

D

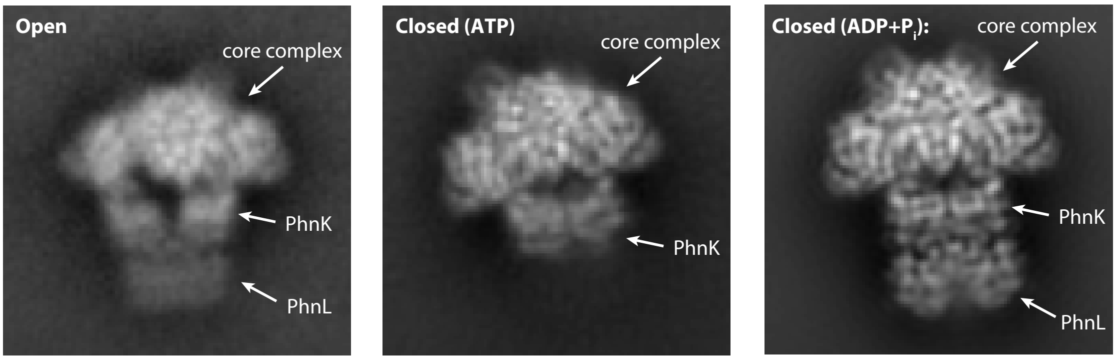

**Supplementary Figure 11**

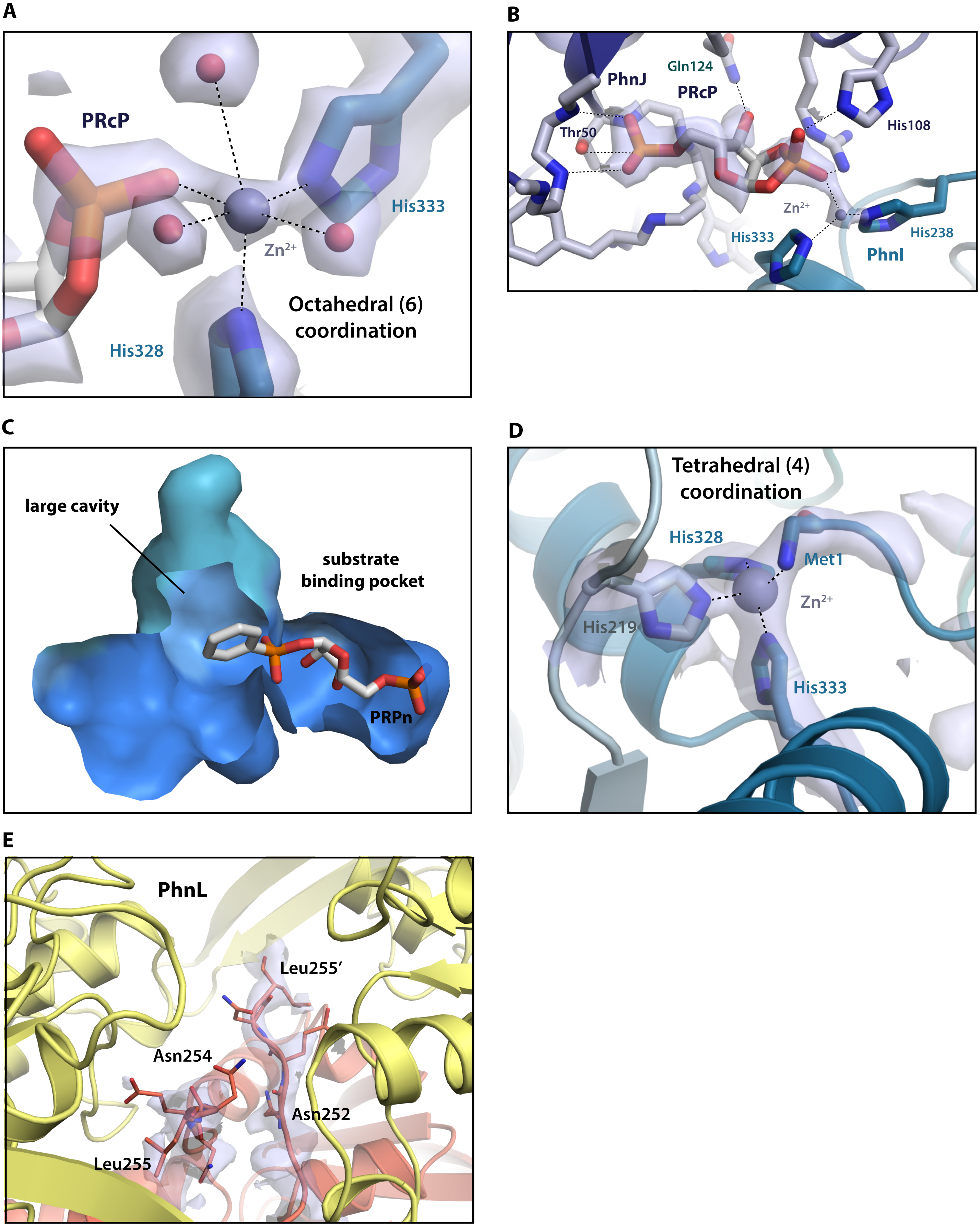

**Supplementary Figure 1 | The C-P lyase pathway.** Pathway for the breakdown of phosphonate compounds by the bacterial C-P lyase pathway. In all intermediates, the phosphorus atom originating from the phosphonate is shown in red and R- indicates a carbon moiety. In step 1, PhnI in the presence of PhnG, PhnH and PhnL catalyses the phosphonylase reaction whereby the adenine base of ATP is exchanged by the phosphonate compound, forming *5-triphosphoribosyl-α-1-phosphonate*. Next, PhnM catalyses the removal of pyrophosphate producing *5-phosphoribosyl-α-1-phosphonate* and $PP_i$ (step 2). *5-phosphoribosyl-α-1-phosphonate* is then the substrate for the *S-adenosylmethionine* (SAM) dependent C-P lyase reaction catalysed by PhnJ (step 3), which liberates the carbon moiety (R-H) and produces *5-phosphoribosyl-1-2-cyclic phosphate*. In step 4, PhnP a phosphoribosyl cyclic phosphodiesterase, hydrolyses the C2 ester resulting in production of *ribose 1,5-bisphosphate*. In step 5, the *ribosyl bisphosphate phosphokinase* PhnN phosphorylates *ribose 1,5-bisphosphate* to PRPP consuming 1 ATP. The last two steps responsible for the formation of $P_i$ are not catalysed by enzymes encoded in the *phn* operon. However, a *phosphoribosyltransferase* can convert a nucleobase (e.g., adenine) and PRPP to a nucleoside 5'-monophosphate (such as AMP) and $PP_i$ (6). Finally, $PP_i$ can then be hydrolysed by a diphosphatase to $P_i$.

**Supplementary Figure 2 | Cryo-EM analysis of Phn(GHIJ)$_2$K.** Diagram showing the overall strategy used to sort picked particles for the Phn(GHIJ)$_2$K structure. Initial steps of 2D and 3D classification in RELION yielding 1,329,400 particles were followed by heterogeneous refinement in cryoSPARC, resulting in 901,800 particles used to generate a 2.2 Å map in RELION. Mask used for 3D classification with signal subtraction and without image alignment are shown as a semi-transparent surface over all maps. The Phn(GHIJ)$_2$ core complex is shown in blue and PhnK in red. Particle classes from 3D classification with signal subtraction and no image alignment used in 3D variability analysis are marked in rounded boxes. The final map generated after 3D variability analysis and homogeneous refinement is shown in the bottom of the figure with accompanying Guinier plot, particle directional distribution plot, FSC curve, and local resolution map.

**Supplementary Figure 3 | Extra density in the ATP-binding site of PhnK.** 3D density maps of PhnK from 3D classification with signal subtraction of the Phn(GHIJ)$_2$K data. In red, the map of a 3D class of PhnK with improved density and in semi-transparent blue a similar map from 3D classification with an extra density at the nucleotide binding site.

**Supplementary Figure 4 | Cryo-EM analysis of Phn(GHIJKL)$_2$ bound to AMPPNP.** Diagram showing the overall strategy used to sort picked particles for the Phn(GHIJKL)$_2$ PhnK E171Q AMPPNP structure. The Phn(GHIJ)$_2$ core complex is shown in blue, PhnK in red, and PhnL in yellow. Initial 3D volumes without interpretable density (Class 3 and 4) are shown in single colours. Particle classes used in the following steps are marked in rounded boxes. The final Phn(GHIJKL)$_2$ PhnK E171Q structure (2.08 Å) was determined by imposing C2 symmetry as shown at the bottom of the figure with accompanying Guinier plot, particle directional distribution plot, FSC curve, and local resolution map.

**Supplementary Figure 5 | Comparison of the ATP binding site in PhnK between different structures. a,** Full map density (top) and details of the PhnK ATP binding site (bottom) with accompanying ligand density of the Phn(GHIJKL)$_2$ PhnK E171Q AMPPNP (left), Phn(GHIJKL)$_2$ WT ADP + Pi (middle), and Phn(GHIJK)$_2$ WT ATP (right) structures. The $Mg^{2+}$ shown in light blue, the catalytic water molecule and other water molecules coordinating the $Mg^{2+}$ are shown as red spheres. The catalytic Glu/Gln171 and accompanying Gln90 residue are shown as sticks. Electron potential density for nucleotide ligands, the $Mg^{2+}$ ion, waters, and visible residues are shown. **b,** Structural alignment of the dimeric ABC modules in the *Staphylococcus aureus* Sau1866 ABC transporter bound to AMPPNP (PDB ID: 2ONJ, green) and PhnK E171Q (red), both bound to

AMPPNP. The nucleotide and residues involved in binding, are shown as coloured sticks. The transmembrane part of Sau1866 is in grey with the coupling helix indicated (cyan). **c,** Superposition of the Phn(GHIJ)$_2$K and Phn(GHIJKL)$_2$ structures shown as ribbons.

**Supplementary Figure 6 | Identification of PhnL and PhnK-PhnL sequence alignment. a,** Top hits from MASCOT following total protein analysis of a purified sample prior ion-exchange chromatography. All the top hits are C-P lyase core complex components (PhnGHIJ), PhnK, or PhnL. **b,** Electrostatic surface potentials of the PhnK (left, top side) and PhnL (right, under side) dimers as observed in the structure of PhnGHIJKL PhnK E171Q AMPPNP shown as a semi-transparent surface on top of cartoon representations. Electrostatic potential scale is from -5.0 to +5.0. **c,** Structural sequence alignment of PhnK, PhnL and *S. aureus* Sav1866 (PDB ID: 2ONJ) with strictly conserved residues marked in green boxes. Secondary structure elements of PhnK and PhnL are shown above the sequences and labelled. The regions containing the extended β hairpin of PhnL and the C-terminal domain are shown in yellow and red boxes, respectively.

**Supplementary Figure 7 | Purification of the Phn(GHIJKL)$_2$ complex and identification of PhnL. a,** Purification of Phn(GHIJKL)$_2$ from pRBS01 expressed in *E. coli* Lemo21 cells. Four discrete peaks identified after MonoQ ion exchange chromatography (Peaks 1-4) were separated and investigated individually by gel filtration chromatography, as indicated. **b**, All peaks contained PhnG, PhnH, PhnI, PhnJ and PhnK, but only Peak 1 contained PhnL as indicated by SDS-PAGE. Peak 1 also contained uncleaved PhnK (PhnK-Strep), despite having been incubated with TEV protease.

**Supplementary Figure 8 | Particle sorting during cryo-EM analysis of PhnGHIJKL under ATP turnover conditions.** Diagram showing the overall strategy used to sort picked particles for the dataset of Phn(GHIJKL)$_2$ under ATP turnover conditions. The Phn(GHIJ)$_2$ core complex is shown in blue, PhnK in red, and PhnL in yellow. Initial 3D volumes without interpretable density (Class 1 and

4) are shown in single colours. Particle classes used in the following steps are marked in black, rounded boxes. The final set of particles used for the reconstruction of the Phn(GHIJKL)$_2$ WT ADP + Pi (1.9 Å), Phn(GHIJK)$_2$ ATP Closed (2 Å), and Phn(GHIJK)$_2$ Open (2.6 Å) structures are shown in orange, red or blue boxes, respectively.

**Supplementary Figure 9 | Final maps from cryo-EM analysis of PhnGHIJKL under ATP turnover conditions.** Final electron potential maps for the Phn(GHIJKL)$_2$ WT ADP+Pi (1.9 Å, top), Phn(GHIJK)$_2$ ATP Closed (2.0 Å, middle), and Phn(GHIJK)$_2$ Open (2.6 Å, bottom) structures with accompanying Guinier plots, particle directional distribution plots, FSC curves, and local resolution maps. The Phn(GHIJKL)$_2$ WT ADP+Pi and Phn(GHIJK)$_2$ ATP closed maps were calculated with imposed C2 symmetry.

**Supplementary Figure 10 | Cryo-EM density features and 2D classes. a,** Details and density at the ATP binding site of PhnL in the Phn(GHIJKL)$_2$ WT structure with density for ATP, $Mg^{2+}$, and Phe11, Gln97, and Glu175 of the A loop shown. **b,** Local resolution maps of the PhnK dimer in the Phn(GHIJKL)$_2$ WT and Phn(GHIJK)$_2$ structures. Both resolution intervals cover 0.8 Å. **c,** 2D-classes of the particles used to generate the Phn(GHIJK)$_2$ Open (top), Phn(GHIJK)$_2$ closed ATP (middle), and Phn(GHIJKL)$_2$ WT (bottom) maps. **d,** Close-up of selected 2D classes showing a side view that reveals the presence or absence of PhnL in the respective particles. Density elements of the C-P lyase Phn(GHIJ)$_2$ core complex, PhnK and PhnL are marked with arrows.

**Supplementary Figure 11 | Ligand binding at the PhnI-PhnJ interface $Zn^{2+}$ site. a,** Details and coordination of the octahedral (6-coordinated) $Zn^{2+}$ at the PhnI-PhnJ binding interface as found in the closed conformation of the Phn(GHIJKL)$_2$ structure. Water molecules are shown as red spheres and coordinating interactions with dashed lines. Density for the $Zn^{2+}$ and all coordinating ligands are shown. **b,** Interactions and density of the 5-phospho-*$\alpha$-D*-ribose-1,2-cyclic-phosphate (PRcP) ligand at the PhnI-PhnJ interface binding site. Protein residues and backbones involved in the interaction with the ligand are shown as sticks and specific interactions are shown with dashed lines. **c,** Binding pocket at the PhnI-PhnJ interface. The substrate binding pocket is shown with 5-phospho-*α*-*D*-ribose-1-phosphonate (PRPn) docked and carrying a phenyl R-group showing that the substrate cavity provides room for larger moieties. **d,** Coordination of the tetrahedral (4-coordinated) $Zn^{2+}$ at the PhnI-PhnJ binding interface in the open conformation of the $Phn(GHIJK)_2$ structure. Density for the $Zn^{2+}$ and all coordinating ligands are shown. **e,** Visible density for the C-terminal PhnK purification tag following the last, natural residue, Asn252, consisting of a TEV protease site, a linker and a 2xStrep purification tag is visible in the density extending from the PhnK C terminus. This extended feature of PhnK is located at the PhnL dimer interface.

**Supplementary Table 1 | Cryo-EM data collection, refinement, and validation statistics**

| | Phn(GHIJ)$_2$K (WT) (EMDB-14445) (PDB 7Z19) | Phn(GHIJKL)$_2$ (PhnK-E171Q) AMPPNP (EMDB-14442) (PDB 7Z16) | Phn(GHIJKL)$_2$ (WT) ADP+P$_i$ (EMDB-14441) (PDB 7Z15) |
|---|---|---|---|
| **Data collection and processing** | | | |
| Magnification | 135,000 | 130,000 | 130,000 |
| Voltage (kV) | 300 | 300 | 300 |
| Electron exposure (e–/Å$^2$) | ~52 | ~62 | ~60 |
| Defocus range (μm) | 0.7-2.0 | 0.5-1.4 | 0.5-1.4 |
| Pixel size (Å) | 0.83 | 0.647 | 0.647 |
| Symmetry imposed | C1 | C2 | C2 |
| Initial particle images (no.) | 901,800 | 1,023,261 | 874,692 |
| Final particle images (no.) | 50,323 | 59,737 | 222,056 |
| Map resolution (Å) | 2.57 | 2.08 | 1.93 |
| FSC threshold | 0.143 | 0.143 | 0.143 |
| Map resolution range (Å) | | | |
| **Refinement** | | | |
| Initial model used (PDB ID) | 4XB6 | 4XB6 | 4XB6 |
| Model resolution (Å) | 2.8 | 2.2 | 2.0 |
| FSC threshold | 0.50 | 0.50 | 0.50 |
| Model resolution range (Å) | | | |
| Map sharpening *B* factor (Å$^2$) | -54.2 | -28.8 | -33.2 |
| Model composition | | | |
| Non-hydrogen atoms | 34,111 | 44,722 | 46,182 |
| Protein residues | 2174 | 2882 | 2888 |
| Ligands | 0 | 2 | 6 |
| *B* factors (Å$^2$) | | | |
| Protein | 84.45 | 35.05 | 21.80 |
| Ligand | - | 35.48 | 27.75 |
| R.m.s. deviations | | | |
| Bond lengths (Å) | 0.005 | 0.003 | 0.011 |
| Bond angles (°) | 0.627 | 0.633 | 0.954 |
| Validation | | | |
| MolProbity score | 1.48 | 1.28 | 1.16 |
| Clashscore | 6.60 | 4.78 | 3.76 |
| Poor rotamers (%) | 0.28 | 1.04 | 0.83 |
| Ramachandran plot | | | |
| Favored (%) | 97.4 | 97.9 | 98.4 |
| Allowed (%) | 2.6 | 2.1 | 1.6 |
| Disallowed (%) | 0.0 | 0.0 | 0.0 |

| | Phn(GHIJK)$_2$ (WT) ATP closed (EMDB-14444) (PDB 7Z18) | Phn(GHIJK)$_2$ (WT) ATP open (EMDB-14443) (PDB 7Z17) |
|---|---|---|
| **Data collection and processing** | | |
| Magnification | 130,000 | 130,000 |
| Voltage (kV) | 300 | 300 |
| Electron exposure (e–/Å$^2$) | ~60 | ~60 |
| Defocus range (μm) | 0.5-1.4 | 0.5-1.4 |
| Pixel size (Å) | 0.647 | 0.647 |
| Symmetry imposed | C2 | C1 |
| Initial particle images (no.) | 874,692 | 874,692 |
| Final particle images (no.) | 81,605 | 31,280 |
| Map resolution (Å) | 1.98 | 2.57 |
| FSC threshold | 0.143 | 0.143 |
| Map resolution range (Å) | | |
| **Refinement** | | |
| Initial model used (PDB ID) | 4XB6 | 4XB6 |
| Model resolution (Å) | 2.0 | 2.7 |
| FSC threshold | 0.50 | 0.50 |
| Model resolution range (Å) | | |
| Map sharpening *B* factor (Å$^2$) | -31.1 | -28.7 |
| Model composition | | |
| Non-hydrogen atoms | 37,705 | 37,614 |
| Protein residues | 2432 | 2432 |
| Ligands | 4 | 1 |
| *B* factors (Å$^2$) | | |
| Protein | 147.68 | 39.82 |
| Ligand | 31.49 | 23.47 |
| R.m.s. deviations | | |
| Bond lengths (Å) | 0.005 | 0.002 |
| Bond angles (°) | 0.870 | 0.467 |
| Validation | | |
| MolProbity score | 1.18 | 1.52 |
| Clashscore | 3.31 | 6.55 |
| Poor rotamers (%) | 1.19 | 0.84 |
| Ramachandran plot | | |
| Favored (%) | 98.1 | 97.1 |
| Allowed (%) | 1.9 | 2.9 |
| Disallowed (%) | 0.0 | 0.0 |
